# Ovo is a master regulator of the piRNA pathway in animal ovarian germ cells

**DOI:** 10.1101/2024.04.23.590802

**Authors:** Azad Alizada, Gregory J Hannon, Benjamin Czech Nicholson

## Abstract

The gene-regulatory mechanisms controlling the expression of the germline PIWI-interacting RNA (piRNA) pathway components within the gonads of metazoan species remain largely unexplored. In contrast to the male germline piRNA pathway, which in mice is known to be activated by the testis-specific transcription factor A-MYB, the nature of the ovary-specific gene-regulatory network driving the female germline piRNA pathway remains a mystery. Here, using *Drosophila* as a model, we combine multiple genomics approaches to reveal the transcription factor Ovo as the master regulator of the germline piRNA pathway in ovaries. The enforced expression of Ovo in somatic cells activates germline piRNA pathway components, including the ping-pong factors Aubergine, Argonaute-3, and Vasa, leading to assembly of peri-nuclear cellular structures resembling nuage bodies of germ cells. Cross-species ChIP-seq and motif analyses demonstrate Ovo binding to genomic CCGTTA motifs within the promoters of germline piRNA pathway genes, suggesting a regulation by Ovo in ovaries analogous to that of A-MYB in testes. Our results also show consistent engagement of the Ovo transcription factor family at ovarian piRNA clusters across metazoan species, reflecting a deep evolutionary conservation of this regulatory paradigm from flies to humans.

## Introduction

The PIWI-interacting RNA (piRNA) pathway is an evolutionary conserved defence system in metazoans that silences transposons in gonads, thereby serving crucial roles in genomic stability, gametogenesis, and fertility ^1–4^. Mutants affecting the piRNA pathway negatively impact the integrity of the germline genome, disrupt gamete development, and typically result in sterility ^5–8^. Host-parasite conflict has driven the piRNA pathway to evolve several specialized adaptations to silence transposons in gonadal cells ^9^. The *Drosophila* ovary is one of the key models that has been used to decipher the workings of the piRNA pathway ^10–18^. Within the *Drosophila* ovary, unlike somatic follicular cells, the germline nurse cells employ a germline-specific version of the pathway encompassing nuage bodies, the ping-pong cycle and non-canonical transcription of dual-strand piRNA clusters ^9,10,19^.

In the nuclei of *Drosophila* germ cells (**Figure 1a**), transcription of piRNA precursors from dual-strand piRNA clusters is driven by Rhino (Rhi), a homolog of the heterochromatin protein 1a (HP1a) ^3,4,20^. Rhi acts in complex with Deadlock (Del) and Moonshiner (Moon), which recruits TBP-related factor 2 (Trf2) to form the transcription initiation complex ^12,18^. Del also interacts with Cutoff (Cuff) to protect the 5′ ends of nascent precursor transcripts ^12,21^. These piRNA precursors are then exported out of the nucleus via a dedicated, non-canonical export machinery utilizing Nuclear Export Factor 3 (Nxf3), Bootlegger (Boot), UAP56 and Nxt1 ^16,22,23^ and are directed to germline-specific, peri-nuclear piRNA processing structures, called nuage, where they are processed into functional piRNAs via the ping-pong cycle ^3,16,22,24,25^.

**Figure 1.**
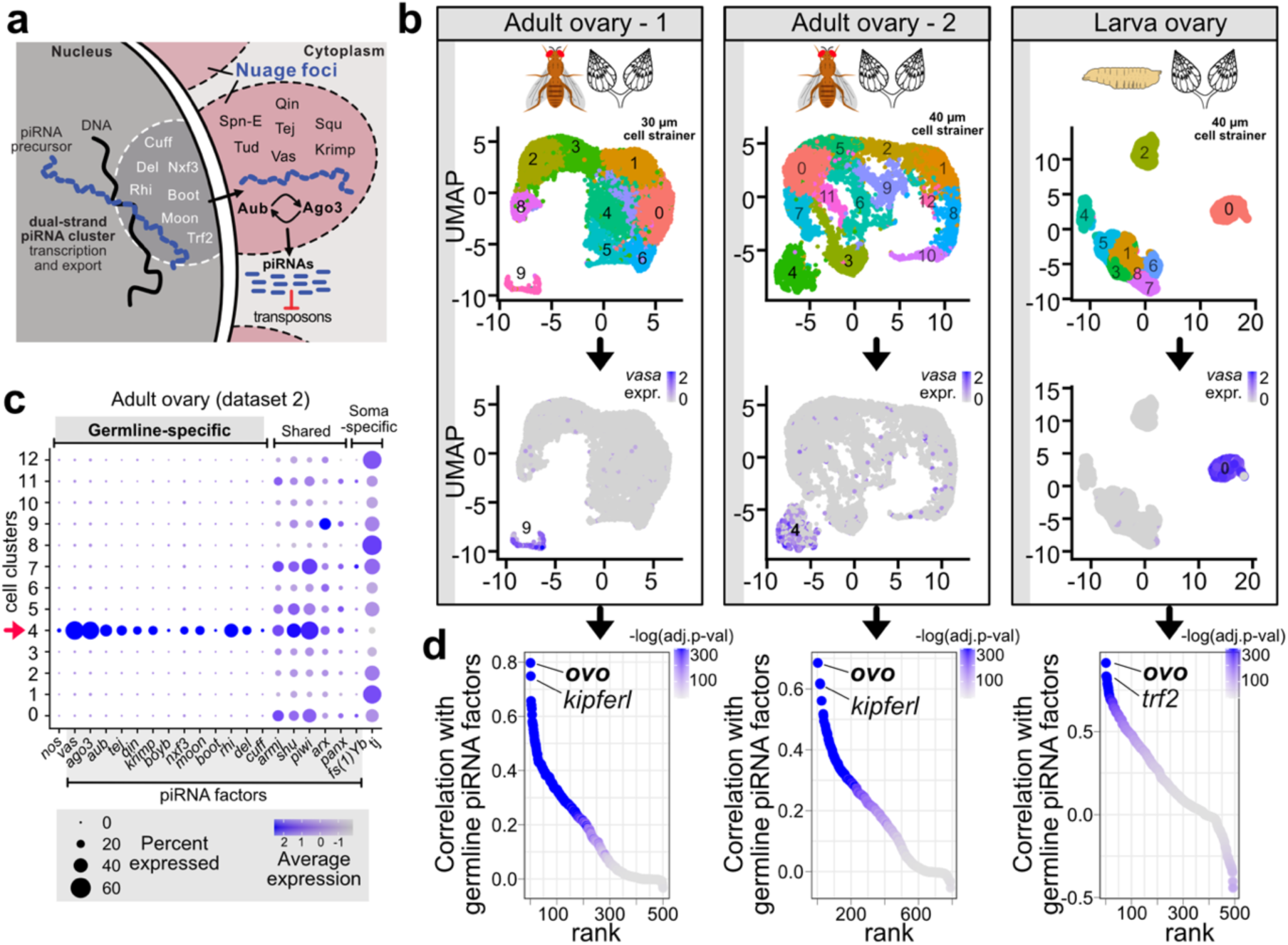
Ovo is the top transcription factor co-expressed with the germline piRNA pathway genes in the *Drosophila* ovary. (a) Model depicting germline piRNA pathway in *Drosophila* germ cells. (b) UMAP clustering of *Drosophila* ovary single-cell RNA-seq datasets (adult ovary 1 from ^35^; adult ovary 2 from ^36^; LL3 larva ovary from ^37^). Expression of the germline marker *vasa* is shown as the marker of the germline clusters. (c) Dot plot showing expressions of the germline-specific, shared, and soma-specific piRNA pathway genes across the clusters identified for the adult ovary 2 dataset (40 µm cell strainer; clusters from the panel a; cluster 4 is the germline cluster). (d) Ranking of DNA-binding proteins and transcription factors by their average expression correlation values (Pearson’s r) with the expression of the germline piRNA pathway factors *aub*, *vas*, and *ago3*. The colour scale indicates correlation p-values adjusted with Bonferroni correction for multiple testing.

In the nuage bodies of *Drosophila* germ cells (**Figure 1a**), ping-pong amplification operates via the PIWI proteins Argonaute-3 (Ago3) and Aubergine (Aub) which cleave piRNA precursor and transposon transcripts in an alternating loop, hence shaping and amplifying the piRNA pool against active transposons ^3,25^. Ago3 in complex with a sense piRNA recognizes and cleaves cluster transcripts through sequence complementarity generating antisense pre-piRNAs that are loaded into Aub ^3,25^. Antisense piRNA-loaded Aub in turn detects and cleaves target transposon mRNAs, thereby forming a new Ago3 complex loaded with a sense piRNA, and completing the cycle ^3,25^. This process is assisted by other nuage-localised protein components, namely, the DEAD-box RNA helicase Vasa (Vas), the putative nuclease Squash (Squ), and Tudor domain proteins Tejas (Tej), Tapas, Qin, Krimper (Krimp) and Spindle-E (Spn-E) ^26–29^. This is in sharp contrast to piRNA biogenesis in somatic cells, which lack nuage bodies and ping-pong amplification, and where piRNA precursors are transcribed from unistrand clusters (e.g., *flamenco*), exported to the cytoplasm by a canonical machinery, and processed into piRNAs via Zucchini-mediated biogenesis^30^.

The gene-regulatory network underlying the control of the germline-specific piRNA pathway in the ovaries of metazoan species have been largely unexplored. The specific expression patterns of piRNA pathway genes in germ cells could be controlled by a positive regulation, such as activation by germline-specific transcription factors (TFs), or via a negative regulation through repression by soma-specific factors. To date, clues from previous studies have pointed towards the existence of both mechanisms.

A study in mouse testes have identified the TF A-MYB as the transcriptional activator of piRNA pathway components during spermatogenesis, including piRNA factors such as *Piwil1* (*Miwi)* and the germline pachytene piRNA clusters ^31^. This study revealed the existence of a positive transcriptional regulator of the male germline piRNA pathway in metazoan testes, yet such transcriptional regulatory pathways controlling the expression of the female germline piRNA pathway in the ovaries of metazoan species, including *Drosophila*, remain to be identified. On the other hand, earlier work has suggested the existence of negative regulation of germline piRNA factors in *Drosophila* somatic cells ^32–34^. Here, germline-specific components of the piRNA pathway such as *aub*, *ago3*, and *vas* were upregulated upon deletion of the tumour suppressor gene *lethal (3) malignant brain tumor* [*l(3)mbt*]. However, the mechanism underlying this regulation remained unclear.

In this study, using *Drosophila* as a model, we uncover key elements of the gene-regulatory network controlling the female germline piRNA pathway. In a systematic analysis integrating multiple approaches, including single-cell RNA-seq, ATAC-seq, ChIP-seq as well as depletion and overexpression screens, we identify the transcription factor Ovo as the key positive transcriptional regulator of the germline piRNA pathway in ovaries. Enforced expression of Ovo in somatic cells activates expression of the germline piRNA pathway genes including the ping-pong cycle components Aub, Ago3 and Vas, leading to formation of peri-nuclear cellular structures mimicking the nuage bodies of germ cells. This is orchestrated through binding of Ovo to conserved CCGTTA motifs within the promoters of these genes. We also reveal the mechanistic link between *l(3)mbt*, *ovo* and the germline piRNA pathway genes. In addition, we show that the ancient Ovo-binding motifs are highly enriched within the germline piRNA clusters in ovaries of metazoan species ranging from insects to humans. ChIP-seq experiments show that fly Ovo and its mouse and human orthologs, OVOL2, are recruited to the motifs within promoters of the piRNA pathway genes and ovarian piRNA clusters. Interestingly, the same CCGTTA motifs are also bound by the male-specific transcription factor A-MYB in testes of vertebrates to control the male germline piRNA pathway ^31^. Our results suggest that Ovo is an ovary-specific counterpart of A-MYB, performing a role in ovaries analogous to that of A-MYB in vertebrate testes. Overall, our results reveal gene-regulatory interactions between the ovary-specific DNA-binding TFs and the ancient CCGTTA *cis*-regulatory elements underlying the control of the germline piRNA pathway in ovaries across the metazoan species.

## Results

### Ovo is co-expressed with germline piRNA pathway genes in *Drosophila* ovary

In a previous study on male piRNA factors in mouse testes, the testis-specific transcription factor A-MYB was identified as a regulator of male germline piRNA pathway genes ^31^. To identify a potential master regulator of the female germline piRNA pathway in the *Drosophila* ovary, we performed a co-expression analysis using three separate single-cell RNA-seq datasets from *Drosophila* ovaries to determine the ovarian TFs co-expressed with the germline-specific piRNA pathway genes, *aub*, *vas*, and *ago3* (**Figure 1b and Supplementary Dataset 1**). Two of the scRNA-seq datasets were from adult ovaries ^35,36^ and one was from larval ovaries ^37^. Clustering of each of the datasets revealed distinct germline clusters showing specific expression of germline markers (e.g., *vas*, *nos*) and germline-specific piRNA pathway genes (i.e., *aub*, *vas*, *ago3*, *tej*, *qin*, *krimp*, *boyb*, *nxf3*, *moon*, *boot*, *rhi*, *del*, and *cuff*) (**Figures 1b and 1c, Supplementary Figures 1a-d**).

Correlation analysis (Pearson’s r) identified *ovo* as the most significantly co-expressed TF with germline-specific piRNA pathway genes in all three datasets (Pearson’s r>0.7 in adult flies and Pearson’s r>0.9 in larva, adjusted p<1.0x10^-300^) (**Figure 1d and Supplementary Dataset 1**). Interestingly, the second most significant co-expressed TF in adult ovaries was *kipferl* (**Figure 1d**), a recently identified zinc-finger protein that serves as a co-factor for the germline piRNA factor, Rhino, which is required for piRNA production from most dual-strand piRNA clusters ^38^. Of note, the second top-ranking TF in larval ovaries was *trf2* (**Figure 1d**), which has been shown to assist Moonshiner with transcription of dual-strand piRNA clusters ^18^.

During *Drosophila* oogenesis, germline stem cells (GSCs) at the tip of germarium continuously generate egg chambers ^39,40^, thus the germ cell cluster identified by scRNA-seq represents an aggregate of stages throughout germline development. To delineate the TFs co-expressed with germline piRNA pathway genes across oogenesis stages, we isolated and re-clustered the germ cell cluster and performed a germline-specific correlation analysis (**Figure 2 and Supplementary Figures 2**).

**Figure 2.**
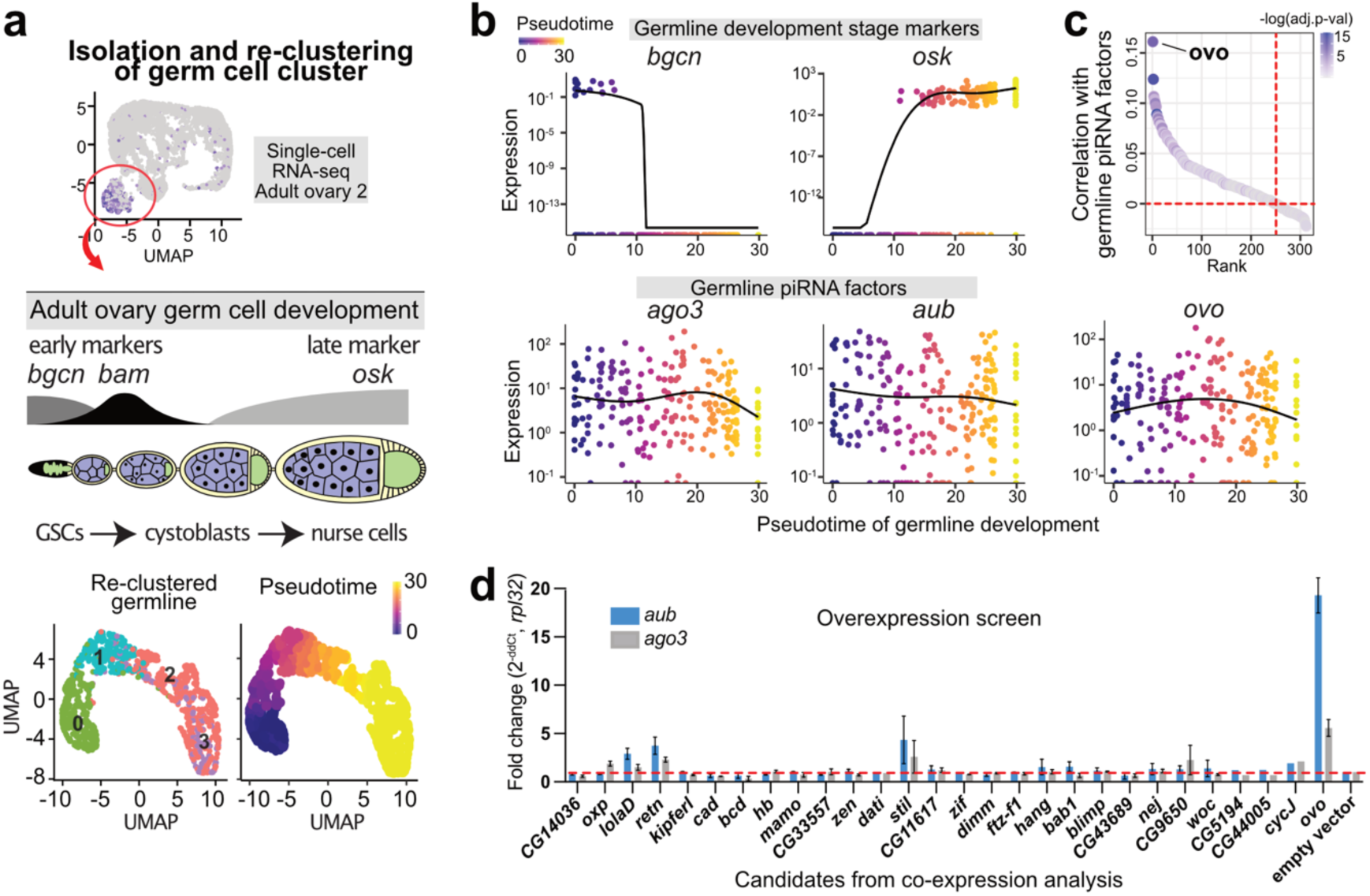
Overexpression screen of the germline co-expressed candidates reveals Ovo as a positive regulator of piRNA pathway genes. (a) Diagram showing isolation and re-clustering of the germ cells (cluster 4) from adult ovary single-cell RNA-seq (dataset 2) and computation of the pseudotime trajectory by rooting the *bgcn*-expressing germline stem cells (GSCs) as the starting point. (b) Expression pattern of the early (*bgcn*) and late (*osk*) stage markers of germline differentiation along the pseudotime trajectory of the germline development shown together with the germline piRNA pathway genes (*ago3* and *aub*) and the top co-expressed transcription factor *ovo*. (c) Ranking of the DNA-binding proteins and transcription factors by the average expression correlation (Pearson’s r) with the germline piRNA pathway genes *aub*, *vas*, *qin*, and *ago3* within the re-clustered germ cell cluster (cluster 4 in adult ovary scRNA-seq dataset 2). The colour scale shows correlation p-values adjusted with Bonferroni correction for multiple testing. (d) Overexpression screen in ovarian somatic cells (OSCs) using the top co-expressed candidates (TFs and chromatin-binding proteins) from the co-expression analysis (RT-qPCR, 48-72hr post-nucleofection in OSCs, n=3 replicates from distinct samples, error bars indicate standard error of the mean). Overexpression of Ovo-B isoform (matching NM_080338) is indicated for *ovo*.

Using early (*bgcn* and *bam*) and late markers (*osk*) of oogenesis ^41–43^, we computed a pseudotime of germline development and tracked the expression pattern of the germline piRNA pathways genes from GSCs to nurse cells (**Figures 2a and 2b, and Supplementary Figures 2a and 2b**). Correlation analysis revealed *ovo* as the top-ranking TF co-expressed with the germline piRNA pathway genes over the course of germline development (adjusted p<3.4x10^-5^, **Figure 2c, Supplementary Figure 2c, and Supplementary Dataset 1**).

To functionally validate the top co-expressed gene-regulatory candidates, we performed an overexpression screen in ovarian somatic cells (OSCs) (**Figure 2d**). Ectopic expression of *ovo* (isoform *ovo-B*, NM_080338 transcript) resulted in upregulation of the germline piRNA pathway genes in OSCs (*aub* ∼20-fold, *ago3* ∼5-fold, *vas* ∼5-fold; p<0.01, RT-qPCR) while other candidates had weaker or no effects on their expression (**Figure 2d**).

### Ovo is the principal germline transcription factor in the *Drosophila* ovary

Differential expression analyses between germline and somatic cell clusters in the three separate ovarian single-cell RNA-seq datasets showed *ovo* as the most enriched germline TF (**Supplementary Figures 3a and Supplementary Dataset 2**). To validate this finding, we next aimed to experimentally identify all germline-enriched TFs within *Drosophila* ovaries that could potentially be responsible for promoting germline fate along with the germline piRNA programme.

To delineate germline-specific TFs, we first performed RNA-seq on FACS-sorted germline (vas-GFP+) and somatic cells (vas-GFP-) from *vas*-GFP *Drosophila* ovaries (**Figure 3a**). A differential RNA-seq analysis between the sorted germline and somatic cells (DESeq2) revealed *ovo* as the top germline-enriched TF (∼87-fold, p<1.0x10^-300^, *ovo-B* isoform matching NM_080338 transcript, **Figure 3a and Supplementary Dataset 2**). Cross-tissue RNA-seq analysis showed ovaries are the most expressed site for *ovo* (**Supplementary Figure 3b**). This result corroborates the hypothesis that *ovo* is the principal ovarian germline TF, potentially controlling a germline-specific expression program encompassing the piRNA pathway in ovarian germ cells. To confirm our findings, we additionally performed RNA-seq on the ovarian germline/somatic co-culture line (fGS/OSS) and compared it to purely somatic ovarian cell line (OSCs), which similarly revealed *ovo* as the most enriched germline-specific TF (∼155-fold, p<1.0x10^-300^, **Figure 3b**).

**Figure 3.**
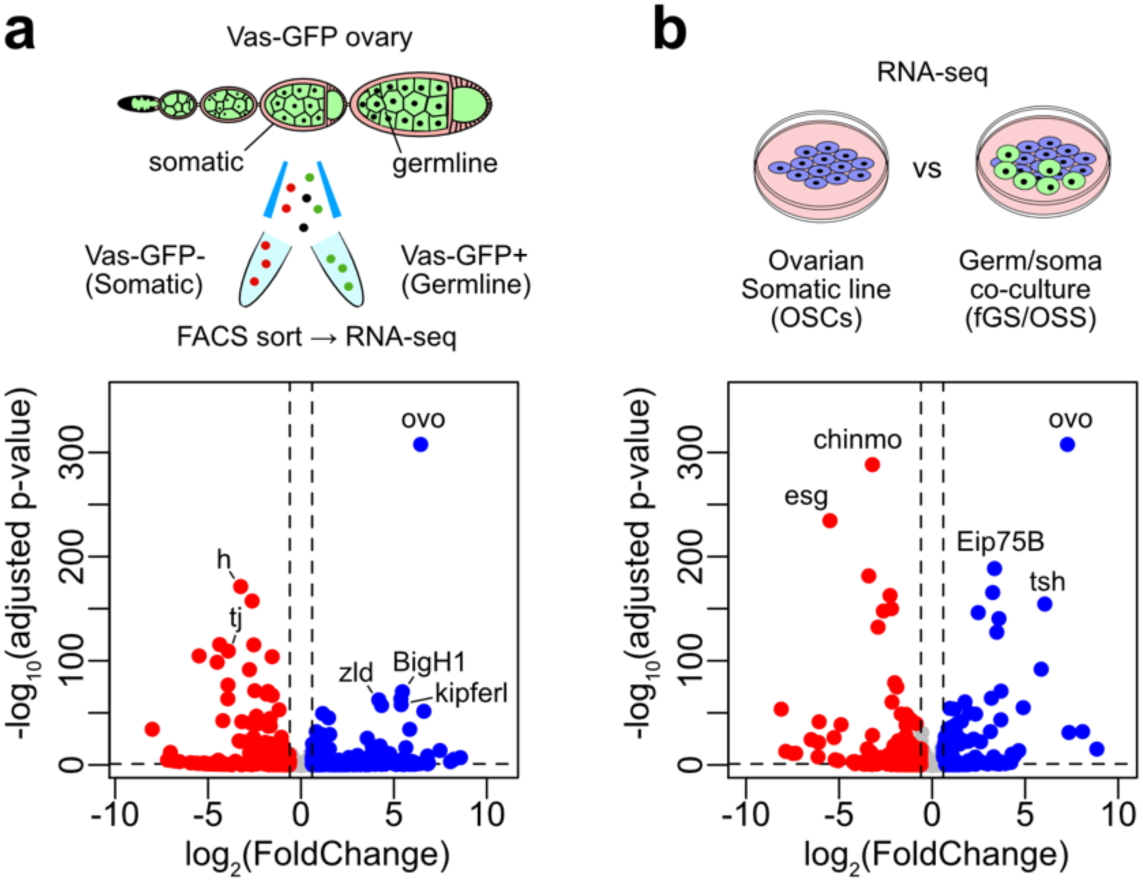
Ovo is the principal transcription factor of the *Drosophila* ovarian germ cells. (a) Differential gene expression of the DNA-binding TFs and chromatin proteins between the FACS-sorted Vas-GFP+ (germline) and Vas-GFP-(somatic) cells from the transgenic Vas-GFP fly ovaries (Deseq2; RNA-seq n=3 replicates from distinct samples). (b) Differential gene expression of the DNA-binding TFs and chromatin proteins between the ovarian somatic cells (OSCs) and the ovarian germ/soma co-culture line (fGS/OSS) (Deseq2; RNA-seq n=4 replicates from distinct samples).

The germline-somatic differential RNA-seq analysis of sorted cells from the *vas*-GFP *Drosophila* ovaries showed a total of 1,019 germline-enriched and 1,316 soma-enriched genes (log_2_FC>1 and log_2_FC<-1, respectively; p.adj<0.01; DESeq2). Among the piRNA pathway genes, only one, *fs(1)Yb*, had a soma-specific expression while 13 components of the germline piRNA pathway, namely, *aub*, *tej*, *ago3*, *vas*, *qin*, *boYb*, *krimp*, *boot*, *moon*, *nxf3*, *cuff*, *rhi* and *del*, showed a strong germline-specific expression (**Supplementary Dataset 2**), and thus could be potentially driven by Ovo.

### Enforced *ovo* expression in ovarian somatic cells activates the germline piRNA pathway components

To determine the gene-regulatory effects of Ovo genome-wide, we performed RNA-seq in ovarian somatic cells (OSCs) following ectopic *ovo* expression (overexpression vector carrying FLAG-tagged *ovo-B* isoform driven by the *act5C* promoter) and compared it to OSCs transfected with an empty vector. Western blotting confirmed the presence of Ovo-B-FLAG protein at the expected size (∼114 kDa) (**Figure 4a**). Immunofluorescence showed that overexpressed Ovo-B-FLAG localized to the nucleus (**Figure 4b and Supplementary Figure 4a**). Differential RNA-seq analysis (DESeq2) showed that ectopic Ovo expression significantly upregulated ∼60-70% of the germline-specific piRNA pathway genes in OSCs, namely, *ago3*, *aub*, *tej*, *qin*, *vas, moon, boot* and *nxf3* (>1.5-fold, adjusted p<0.05, **Figure 4c and Supplementary Figures 4b and 4c and Supplementary Dataset 2**). The strongest effects were seen on *aub* and *ago3* (16-fold and 6-fold, adjusted p<1.0x10^-41^, **Figure 4c and Supplementary Figures 4b and 4c**). In contrast to components of the germline piRNA pathway, the soma-specific *fs(1)Yb* was downregulated upon Ovo overexpression, with all other somatic factors involved in biogenesis and transcriptional gene silencing (TGS) not changing in expression (**Figure 4c and Supplementary Figure 4c**).

**Figure 4.**
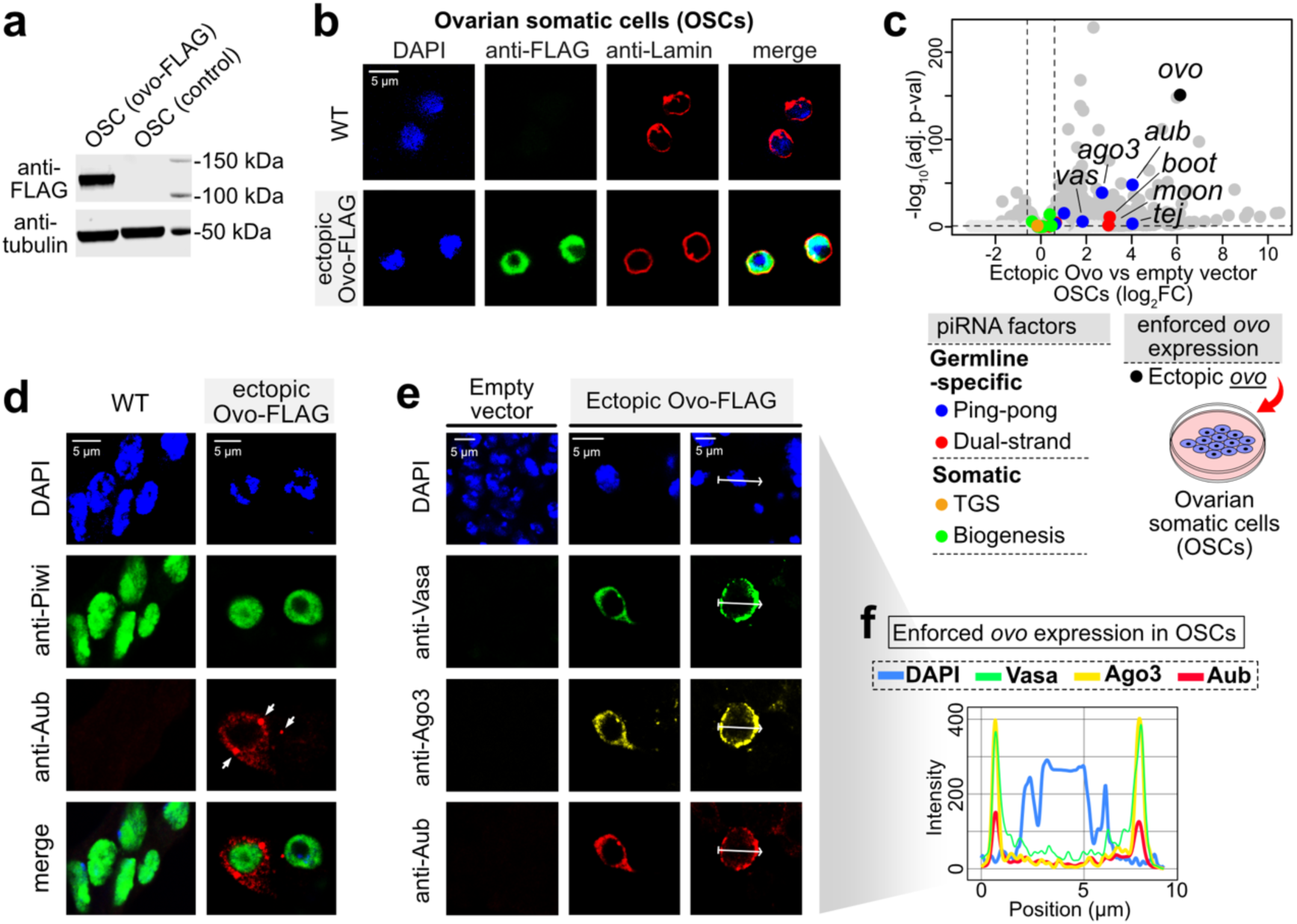
Enforced expression of *ovo* in ovarian somatic cells (OSCs) activates the germline piRNA pathway components leading to formation of cellular structures resembling nuage bodies of germ cells. (a) Western blot showing the presence of the Ovo-FLAG protein following ectopic expression in OSCs. (b) Immunofluorescence images showing the nuclear localization of the Ovo-FLAG protein in OSCs after nucleofection with the *ovo-FLAG* construct (48 hr; Ovo-B isoform, NM_080338 transcript). DAPI indicates DNA and Lamin indicates the nuclear envelope. (c) Differential gene expression between *ovo-FLAG* nucleofected OSCs relative to empty vector (Deseq2; RNA-seq n=3 replicates from distinct samples; the piRNA pathway genes labelled according to the colour key, TGS =transcriptional gene silencing). (d-e) Immunofluorescence images showing the appearance of the nuage components, Aub, Ago3, and Vas, as peri-nuclear nuage-like structures and foci (arrowheads in d) in the *ovo-FLAG* nucleofected OSCs (Piwi indicates the nucleoplasm, DAPI indicates DNA). (f) Colocalization of Aub, Ago3 and Vas proteins within the nuage-like bodies formed around the nuclei (DAPI) of the *ovo-FLAG* nucleofected OSCs. The fluorescence intensity along the white arrow inset is normalized to the highest value.

Using immunofluorescence, we observed the appearance of Aub, Ago3, and Vas proteins following ectopic Ovo expression in OSCs which assembled into peri-nuclear “nuage”-like structures (**Figures 4d-f and Supplementary Figure 4d**). Co-localization analysis showed that Aub, Ago3, and Vas proteins assembled as foci around nuclei in a ring-shaped manner (**Figures 4e and 4f**), resembling the nuage structures of the *Drosophila* germ cells where piRNAs are processed by the ping-pong cycle. Of note, we confirmed that these “nuage”-like structures were distinct from the somatic Yb-bodies of OSCs (**Supplementary Figure 4e**).

A total of 693 genes were upregulated by ectopic Ovo expression in OSCs (log_2_FC>0.6, p.adj<0.1; DESeq2), 182 of which (∼26%) showed strong germline-enriched expression in ovaries (*vas*-GFP log_2_FC>1, p.adj<0.01) (**Supplementary Dataset 2**). Ovo could account for the regulation of at least ∼18% of all germline-enriched genes in ovaries (182 out of 1,019 genes). Notably, among the top germline targets of Ovo was *nanos* (*nos*) (**Supplementary Figure 4f**), a gene essential for germline formation and maintenance ^44,45^. Overall, our results show that Ovo upregulates the majority of the germline-specific piRNA pathway genes when ectopically overexpressed in OSCs. Next, we sought to decipher mechanisms of Ovo regulation to understand how it activates piRNA factor expression exclusively in germ cells.

### L(3)mbt suppresses the germline piRNA pathway by repressing *ovo* in ovarian somatic cells

Early clues regarding the regulation of germline piRNA pathway components in *Drosophila* came from studies that deleted the *l(3)mbt* gene in somatic tissue, including the brain, ovary and ovary-derived OSCs, which uniformly resulted in the upregulation of several germline-specific genes such as the ping-pong components Aub, Vas, and Ago3 ^32–34^. The molecular mechanism underlying this activation has remained unknown. L(3)mbt has been shown to interact with chromatin and cause histone compaction leading to suppression and insulation of gene expression ^46–48^. While a direct regulation by L(3)mbt could explain prior observations, we hypothesized that the deletion of L(3)mbt could also upregulate one or several specific TFs that in turn control the expression of germline piRNA pathway genes.

To determine the precise mechanistic link between L(3)mbt and germline piRNA pathway components, we performed RNA-seq (n=3 replicates) and ATAC-seq (n=2 replicates) on wild-type (WT) OSCs and OSCs carrying a deletion of *l(3)mbt* [Δ*l(3)mbt*] (**Figure 5a**). We aimed to uncover regulatory elements, transcription factors, and genes that are differentially regulated and could be responsible for the activation of the germline piRNA pathway components upon L(3)mbt deletion.

**Figure 5.**
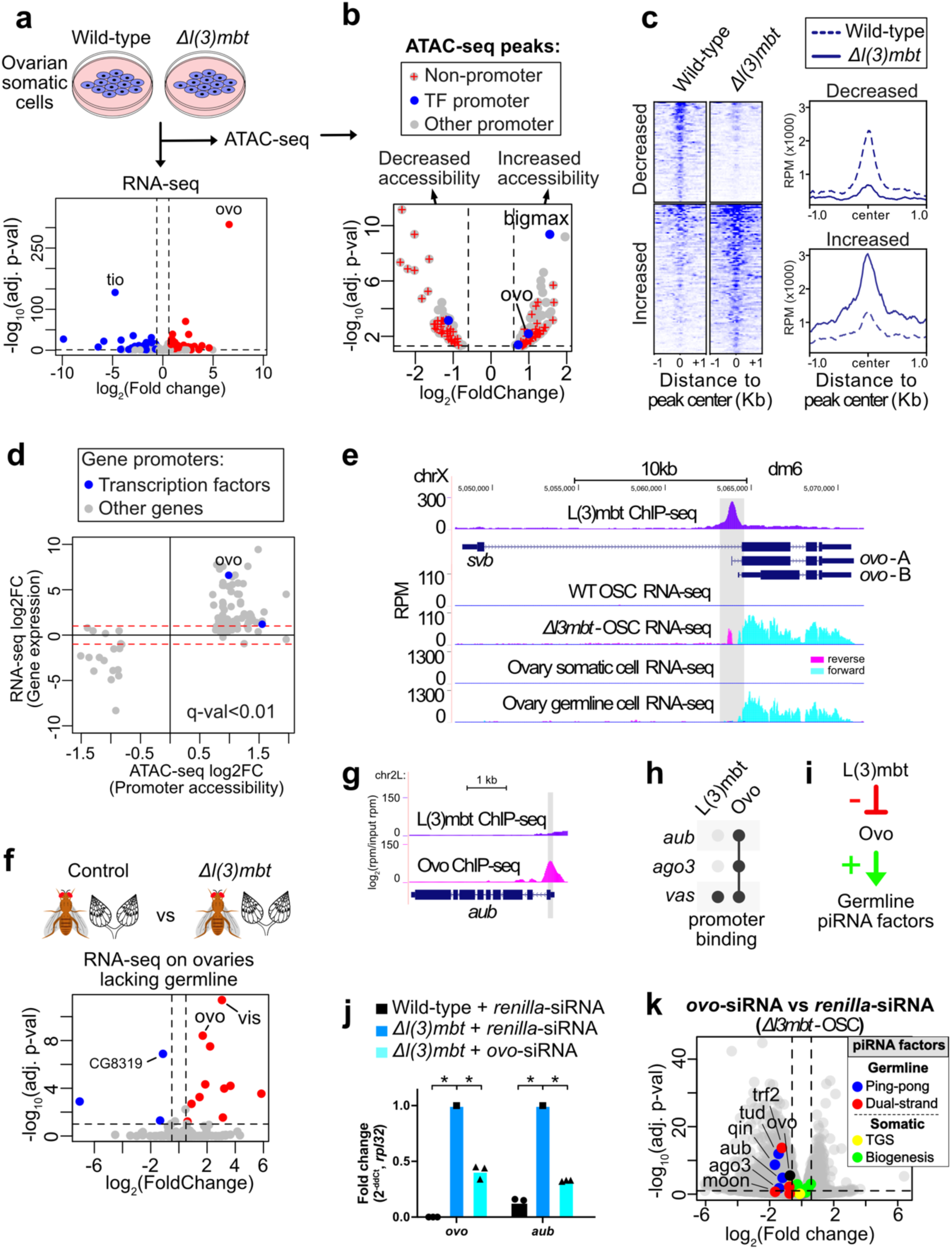
L(3)mbt negatively regulates the germline piRNA pathway via repression of *ovo* expression in somatic cells. (a) Differential gene expression of the DNA-binding transcription factors (TF) between the wild-type (WT) ovarian somatic cells (OSCs) and Δ*l(3)mbt* OSCs (Deseq2; RNA-seq n=4 replicates from distinct samples). (b) Differential chromatin accessibility between the WT OSCs and Δ*l(3)mbt* OSCs (DiffBind; ATAC-seq; n=2 replicates from distinct samples). Gene promoters are defined as ATAC-seq peaks overlapping ±1kb region of gene transcription start site (TSS). (c) Heatmaps and genomic profiles of ATAC-seq reads corresponding to regions of increased and decreased chromatin accessibility (FDR q <0.05; DiffBind). (d) All genes showing significant differential gene expression (RNA-seq; Deseq2, adjusted p-value <0.01) and promoter chromatin accessibility (ATAC-seq; 1kb ± of gene TSS; DiffBind, FDR q <0.01) between WT and Δ*l(3)mbt* OSCs. (e) L(3)mbt ChIP-seq from OSCs showing L(3)mbt binding to the promoter of the *ovo* gene (n=2 replicates from distinct samples; merged; data from ^49^). RNA-seq tracks below showing the expression of the germline *ovo* isoform in Δ*l(3)mbt* OSCs and the *Drosophila* ovary germline cells (n=4 replicates from distinct samples; merged). (f) Differential gene expression (Deseq2) between the control and Δ*l(3)mbt* fly ovaries, both lacking germline due to *tud* maternal mutations (*tud^M^*) (n=3 replicates from distinct samples; RNA-seq data from ^33^). (g) L(3)mbt ChIP-seq from OSCs showing absence of L(3)mbt binding at the germline *aub* promoter (n=2 replicates from distinct samples; merged; data from ^49^). Ovo ChIP-seq showing Ovo binding to the *aub* promoter (n=2 replicates from distinct samples; merged; data from ENCODE, whole-fly) (h) L(3)mbt and Ovo ChIP-seq peaks within the promoters of the ping-pong genes *aub*, *ago3*, and *vas* (±1kb of TSS). (i) Model for the L(3)mbt and Ovo regulation of the germline piRNA pathway genes. (j) Ovo siRNA knockdown experiments in *Δl(3)mbt*-OSCs (RT-qPCR; n=3 replicates from distinct samples; p-value: *<0.01, one-tailed two-sample t-test). (k) Volcano plot showing downregulation of the germline-specific piRNA pathway genes on day 2 of *ovo* siRNA knockdowns in *Δl(3)mbt*-OSCs using differential RNA-seq analysis (Deseq2) between *ovo* and *renilla* siRNA knockdowns (n=3 replicates from distinct samples).

Differential expression analysis of DNA-binding TFs (DESeq2 on RNA-seq data) between Δ*l(3)mbt* and wild-type OSCs revealed *ovo* as the top upregulated TF upon *l(3)mbt* deletion (∼97-fold, adjusted p value <1.0x10^-300^) (**Figure 5a**). The degree of *ovo* upregulation was markedly higher compared to other TFs (**Figure 5a**). To assess if this expression difference was due to increased accessibility at the *ovo* promoter upon *l(3)mbt* loss, we analysed differential chromatin accessibility using ATAC-seq data (DiffBind) from Δ*l(3)mbt* and wild-type OSCs (**Figures 5b-d**). This analysis uncovered 207 genomic regions with significantly increased accessibility upon *l(3)mbt* deletion (>1.5-fold, FDR q <0.05) of which 111 were at gene promoters (±1kb of TSS) (**Figures 5b and 5c**). Out of these, 30 were germline-specific and included only one germline TF, *ovo* (∼2-fold, FDR q <0.01) (**Figures 5d**). The only other TF that showed increased promoter accessibility in Δ*l(3)mbt*-OSCs was *bigmax*, which was not germline-enriched in ovaries. Among the piRNA pathway genes, only *vas* showed increased promoter accessibility in Δ*l(3)mbt*-OSCs and thus could potentially be a piRNA factor that is directly regulated by L(3)mbt, with others likely controlled via Ovo.

We speculated that direct binding of L(3)mbt to the *ovo* promoter could be responsible for the decreased chromatin accessibility in somatic cells, leading to its soma-specific transcriptional repression. Indeed, ChIP-seq data from OSCs showed strong L(3)mbt binding specifically at the promoter region of *ovo* (**Figure 5e**), which increased in accessibility upon *l(3)mbt* deletion. This suggests that direct L(3)mbt binding to the *ovo* promoter in somatic cells represses its expression. Elimination of *l(3)mbt* consequently leads to increased accessibility at the *ovo* promoter and results in drastically increased expression of *ovo*, mimicking its germline expression in the *Drosophila* ovary (**Figure 5e**). We further confirmed our results by reanalysing recently published RNA-seq data from *l(3)mbt* knockdown in OSCs ^49^ which also showed *ovo* as the most significantly upregulated gene (**Supplementary Figure 5a**).

To extend our findings to an *in vivo* setting, we analysed previously published RNA-seq data from Lehmann and colleagues from *l(3)mbt* mutant and control ovaries of transgenic flies that lack germ cells due to maternal *tud* mutations (*tud^M^*) ^33^. This allowed us to perform differential expression analysis between Δ*l(3)mbt* and control ovaries that were devoid of germline cells (**Figure 5f**). This analysis identified *ovo* as the second most significantly upregulated TF upon *l(3)mbt* deletion *in vivo* within somatic cells of fly ovaries (**Figure 5f**). It was previously shown that L(3)mbt forms a complex with Lint-O at chromatin to silence gene expression ^49^. Differential expression analysis of the RNA-seq data from *lint-O* and control knockdowns in OSCs, revealed *ovo* as the top upregulated TF upon Lint-O depletion (**Supplementary Figure 5b**), suggesting that Lint-O together with L(3)mbt forms a repressive complex at the *ovo* promoter in somatic cells, which we confirmed using Lint-O ChIP-seq data in OSCs (**Supplementary Figure 5c**).

We then analysed if L(3)mbt could be exerting its effects on germline piRNA pathway genes, such as *aub,* indirectly via regulation of Ovo. Most importantly, using Ovo ChIP-seq, we observed strong Ovo binding at the promoters of germline-specific piRNA pathway genes where L(3)mbt binding was lacking (**Figures 5g and 5h**), suggesting that L(3)mbt was not directly controlling their expression. We therefore hypothesized that L(3)mbt indirectly represses the expression of germline-specific piRNA pathway genes in somatic cells via its regulation of *ovo* expression (**Figure 5i**).

To establish a causal link, we performed siRNA knockdowns of *ovo* in Δ*l(3)mbt* OSCs, which express *ovo* due to absence of L(3)mbt repression (**Figure 5j**). Knockdown of *ovo* in Δ*l(3)mbt* OSCs resulted in the downregulation of germline piRNA pathway genes, including *aub* and *ago3* (p<0.01, RT-qPCR), further supporting our hypothesis (**Figure 5j and Supplementary Figure 5d**). To validate our finding, we additionally performed RNA-seq on *ovo* siRNA knockdowns in Δ*l(3)mbt* OSCs (**Figure 5k and Supplementary Figure 5e**), which confirmed decreased expression of germline piRNA pathway genes, including *aub* (∼2.3-fold decrease, p.adj=2.8x1.0^-6^; DESeq2), *qin* (∼3-fold decrease, p.adj=2.6x1.0^-8^), *ago3* (∼3-fold decrease, p.adj=0.01) (**Figure 5k and Supplementary Figure 5e**).

Overall, in our analysis we observed that ∼26% (n=315) of all upregulated genes in Δ*l(3)mbt* OSCs (n=1,195) were germline-enriched, representing ∼16% of all germline-enriched genes (n=1,911; log_2_FC>0.6, p.adj<0.1). These genes could be either under a direct L(3)mbt repression (i.e., L(3)mbt promoter binding) or under an indirect repression via germline TFs such as Ovo which are repressed by L(3)mbt (**Supplementary Dataset 2**). Of the 315 L(3)mbt-repressed germline genes, 103 were upregulated by ectopic *ovo* expression alone in OSCs; therefore, ∼33% of the upregulated germline genes in Δ*l(3)mbt* OSCs were likely to be indirectly regulated through activation of *ovo* expression in the absence of *l(3)mbt* (**Supplementary Dataset 2**). RNA-seq of *ovo* siRNA knockdowns in Δ*l(3)mbt* OSCs revealed that ∼35% (n=419 genes) of all L(3)mbt-repressed genes in OSCs (n=1,195) were downregulated by day 2 of *ovo* siRNA knockdown and thus could be accounted for by indirect regulation via Ovo (with approximately 27%, 112 out of the 419 genes, being germline-enriched; **Supplementary Dataset 2**). Out of a total of 2,395 downregulated genes by day 2 of *ovo* siRNA treatment, 488 were germline-enriched including the germline piRNA pathway genes. Thus, we could conclude that at least ∼20% of all upregulated germline genes in Δ*l(3)mbt* OSCs were upregulated indirectly due to activation of Ovo. This fraction remained at ∼20% (485 out of 2,394) by day 4 of *ovo* siRNA treatment (second nucleofection on day 2; **Supplementary Dataset 2 and Supplementary Figure 5e**).

### Ovo regulates the germline piRNA pathway genes via binding to conserved CNGTTA motifs

To capture Ovo binding events that lead to activation of the germline piRNA pathway genes, we performed Ovo ChIP-seq (n=2) following enforced *ovo* expression in OSCs (n=4769 peaks, FDR<0.05, >5-fold enrichment). Using Ovo ChIP-seq data from adult female flies (ENCODE ^50^) we were also able to map *in vivo* Ovo binding events (n=4,477 peaks) across the *Drosophila* genome (**Figure 6a**). We performed ATAC-seq in *Drosophila* ovaries to locate Ovo binding regions corresponding to open chromatin (n=4,007) and closed chromatin (n=470) within ovaries (**Figure 6a**). Ectopic Ovo expression in OSCs recapitulated ∼65% of the *in vivo* binding events observed in flies of which the majority (∼93%) were within open chromatin in ovaries (**Figure 6a**).

**Figure 6.**
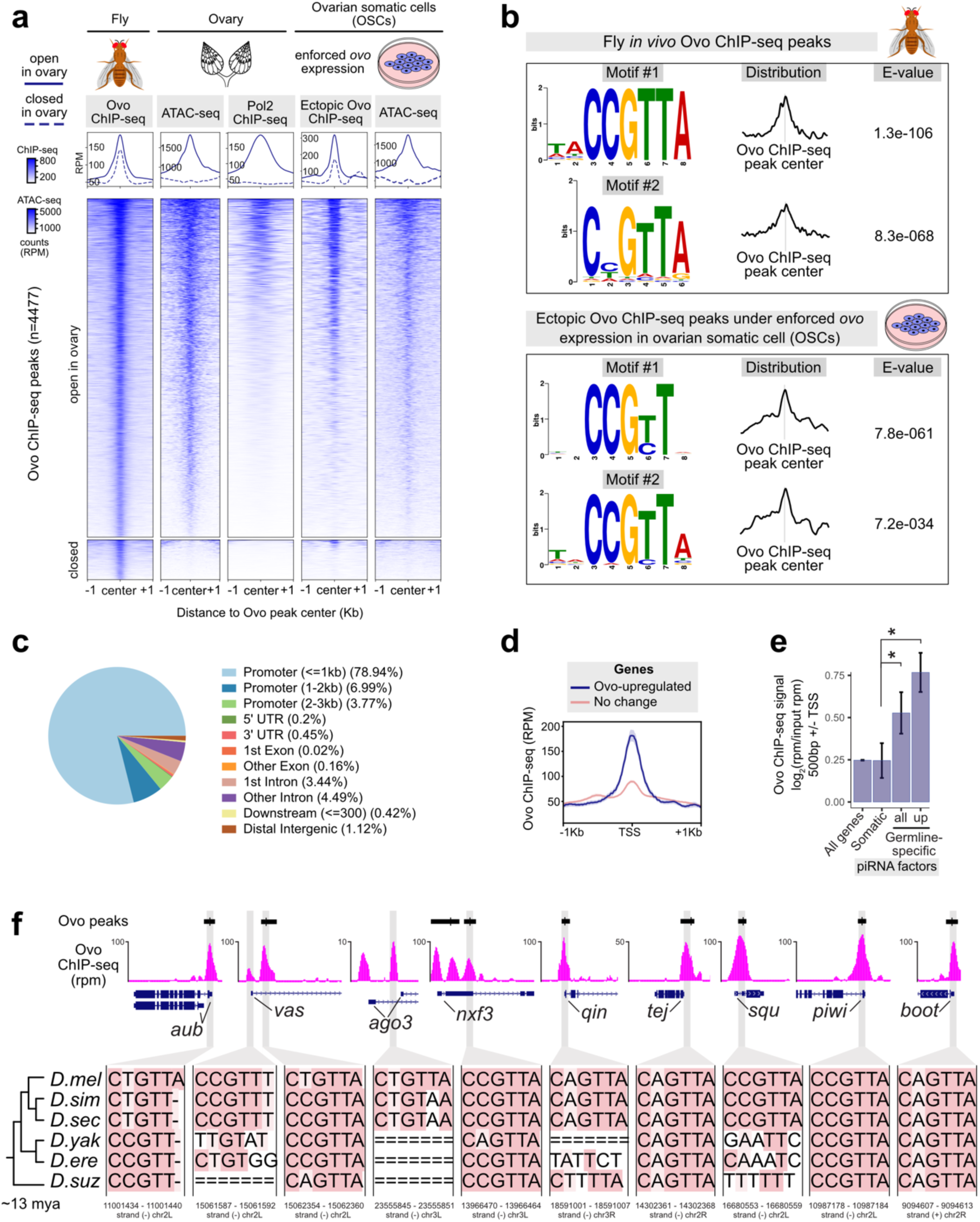
Ovo regulates germline piRNA pathway genes via binding to conserved CNGTTA motifs. (a) Heatmap of fly Ovo ChIP-seq peaks (n=2 replicates from distinct samples merged; ENCODE) corresponding to open and closed chromatin states in fly ovaries based on ovary ATAC- seq peaks (n=2 replicates from distinct samples merged). The binding pattern of the ectopic Ovo ChIP-seq following enforced *ovo* expression in OSCs (n=2 replicates from distinct samples) and ATAC-seq accessibility of OSCs (n=2 replicates from distinct samples) centred on the fly Ovo ChIP-seq peaks are shown by the heatmaps on the right. The engagement of RNA polymerase II (Pol2) in ovaries is also depicted using the ovary Pol2 ChIP-seq signals (n=1; data from ^18^). (b) The top-scoring *de novo* motifs discovered within Ovo ChIP-seq peaks using MEME-ChIP. (c) Genomic annotations (UCSC; Ensembl genes; dm6) of the fly Ovo ChIP-seq peaks using ChIPseeker. (d) Genomic profile of fly Ovo ChIP-seq signals (rpm) within 1kb ± of TSS of the upregulated, downregulated, and unresponsive genes following enforced *ovo* expression in OSCs is shown. (e) Bar plots comparing fly Ovo ChIP-seq signals (input-normalized; rpm) between the promoters of the germline-specific and the somatic piRNA pathway genes. (f) The conservation of Ovo motifs within Ovo ChIP-seq peak summits at the germline piRNA pathway gene promoters is shown by multiple sequence alignments across six *Drosophila* species (27-way insect Multiz alignments and the phylogenetic tree from the UCSC genome browser conservation tracks, Ovo motif coordinates for dm6 genome are shown below).

The top scoring *de novo* motif within *in vivo* Ovo ChIP-seq peaks in flies was C<u>C</u>GTTA (MEME-ChIP, e-value=1.3^-106^), while the secondary motif was C<u>N</u>GTTA (MEME-ChIP, e-value=8.3^-68^) (**Figure 6b**). Interestingly, the C<u>C</u>GTT core of the motif was the top hit within the ectopic Ovo ChIP-seq peaks following enforced *ovo* expression in OSCs (MEME-ChIP, e-value=7.8^-061^) (**Figure 6b**) and C<u>N</u>GTT was the fourth strongest hit *in vivo* (**Supplementary Figures 6c**). These results suggest that C<u>C</u>GTTA is the most preferred Ovo binding site *in vivo*, with changes to the second and last nucleotides often tolerated. The majority of Ovo binding events (∼79%) were proximal to promoters (<1kb to the TSS), while only 1.12% were distal or intergenic (**Figure 6c**). ∼95% of Ovo binding events at promoter proximal regions corresponded to open chromatin in ovaries (based on ATAC-seq peaks) (**Supplementary Figures 6a and 6b**). The genes upregulated in response to enforced *ovo* expression in OSCs showed >2-fold stronger Ovo ChIP-seq binding signals and peak enrichments in their promoters (±1kb of TSS, p<0.01) compared to the genes that did not change in expression (**Figure 6d and Supplementary Figure 6d**). Correspondingly, higher numbers of Ovo motifs were observed at the upregulated Ovo target gene promoters when compared to genes that did not change in expression (∼1.6-fold, p<0.01; **Supplementary Figures 6e-f**).

We then looked at Ovo binding events within promoters of the piRNA pathway genes (**Figures 6e and 6f**). The average Ovo binding signal (input-normalized ChIP-seq) was >2-fold stronger (p<0.05) at promoters of the germline-specific piRNA pathway genes when compared to somatic or shared piRNA factors (**Figures 6e**). This was generally true for all germline genes, which showed on average ∼2-fold stronger Ovo ChIP-seq signal (p<4.1x1.0^-8^) at their promoters (±0.5kb of TSS) when compared to the somatic or shared genes. Moreover, ∼90% of the germline-specific piRNA pathway genes had an Ovo motif ±0.5kb of their TSS overlapping an Ovo ChIP-seq peak summit while only ∼30% of somatic factors did (**Supplementary Figure 4c**). Multiple sequence alignments of the Ovo binding sites within promoters of the germline piRNA pathway genes showed high conservation of the CNGTTA motifs across six *Drosophila* species (*D. melanogaster*, *D. simulans*, *D. sechellia*, *D. yakuba*, D. erecta, *D. suzuki*), sharing a common ancestor about ∼13 million years ago ^51^, suggesting an evolutionary conserved gene-regulatory mechanism behind Ovo interaction with CNGTTA motifs (**Figure 6f**). These results indicate that direct binding to CNGTTA motifs is a major conserved mechanism through which Ovo controls expression of the germline piRNA pathway genes in the *Drosophila* species.

### Vertebrate homologs of Ovo bind to ovarian piRNA pathway components

Most vertebrates have three homologs of the fly *ovo* gene (e.g., mouse *Ovol1*, *Ovol2*, and *Ovol3*) with the *Ovol2* paralog involved in the development of primordial germ cells (PGCs) ^52,53^. Clustering of all vertebrate TF motifs in the JASPAR database (2,022 vertebrates CORE) revealed high similarities between the motifs of Ovo family TFs and, remarkably, A-MYB (**Figure 7a**), a testis-specific TF previously reported to control transcription of both the germline piRNA pathway genes and piRNA clusters in male mice. We hypothesized that Ovo homologs could provide the ovary-specific equivalent of this regulation in females, controlling germline piRNA clusters in ovaries in addition to piRNA pathway genes via the same CCGTTA motifs.

**Figure 7.**
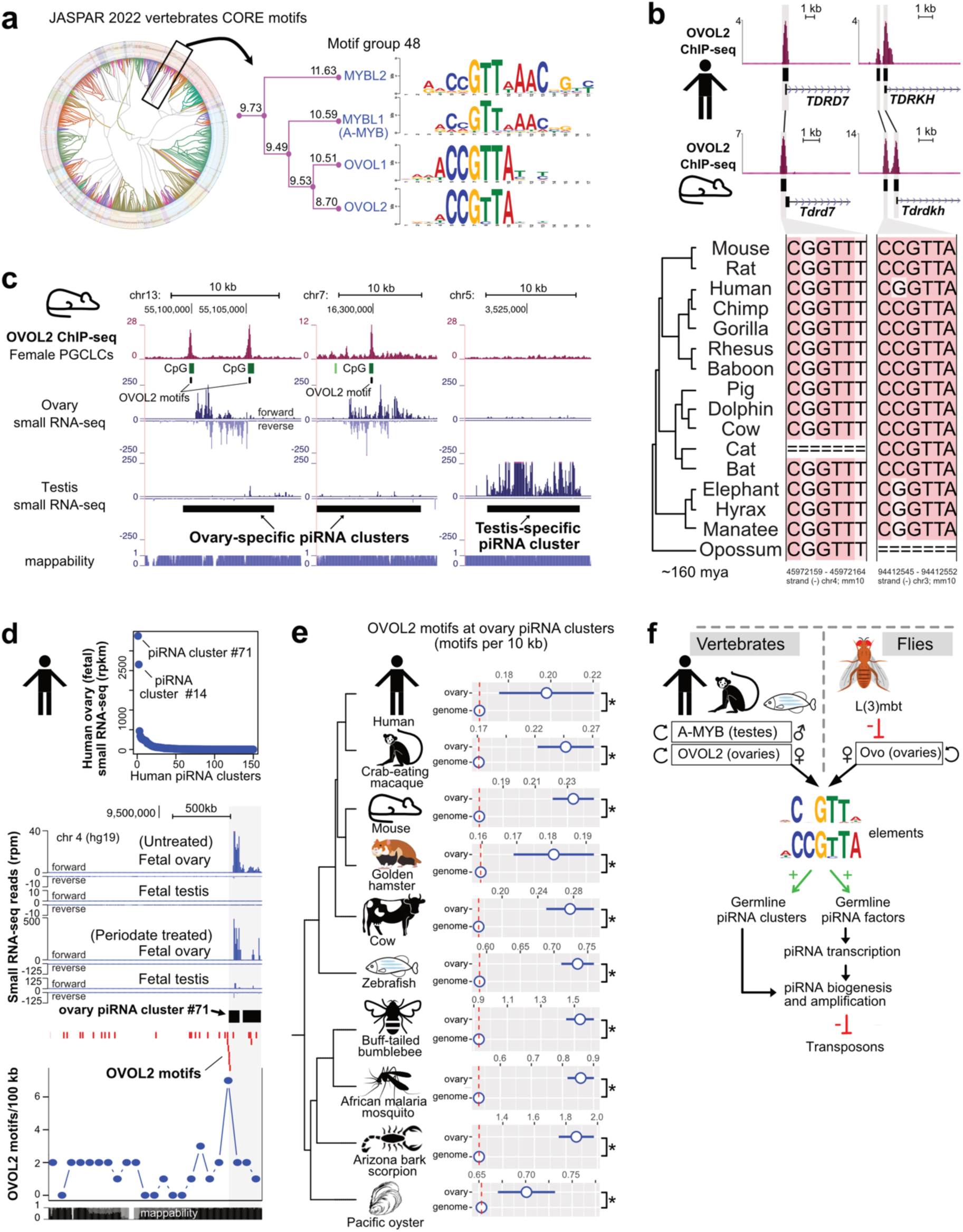
Ovo binding sites are hallmarks of ovarian piRNA pathway components across the genomes of metazoan species. (a) Radial tree clustering of all human TF motifs in the JASPAR database (JASPAR 2022 vertebrates CORE; RSAT -matrix-clustering) showing a zoomed-in view of the motif group 48 containing A-MYB and OVOL2 motifs. Information content of each branch is taken from the dynamic logo forest in JASPAR matrix clustering. (b) Human and mouse OVOL2 ChIP-seq showing OVOL2 binding events at orthologous genomic regions corresponding to promoters of the piRNA pathway genes *TDRD7* and *TDRKH* (rpm; merged n=2 replicates from distinct samples; mouse data is from day 2 of female mouse primordial germ cell-like cell (PGCLC) induction overexpressing mouse *Ovol2a* ^52^ ; human data is from iPSC line overexpressing human OVOL2 ^57^). Multiple sequence alignments of OVOL2 motifs at peak summits indicate high degree of conservation of Ovo binding sites across mammalian species (Multiz alignments of 46 vertebrates from UCSC conservation track, OVOL2 motif coordinates for mm10 are shown below). (c) OVOL2 ChIP-seq from female mouse PGCLCs (day 2 of PGCLC induction; overexpressing transgenic mouse *Ovol2a*; data from ^52^) showing OVOL2 binding to the CCGTTA motifs (OVOL2 motifs) within the mouse ovary-specific piRNA clusters (small RNA-seq data from ^54^). (d) Ranking of the human piRNA clusters by their expression levels in human fetal ovaries (rpkm; data from ^56^). The genomic browser below depicting OVOL2 motifs (red; UCSC squish track display) and the number of OVOL2 motifs per 100 kb at the top-expressed human ovary piRNA cluster (#71). Human fetal ovary and fetal testis small RNA-seq signals show the relative production of the piRNAs from the cluster #71 (rpm; data from ^56^). Sodium periodate treatment enriches for piRNAs. (e) Numbers of OVOL2 motifs per 10 kb compared to the genomic backgrounds across ten metazoan species. Ovarian piRNA clusters were identified using ovary small RNA-seq data from each species (**Supplementary Dataset 3**). The classification tree is based on the NCBI taxonomy database. Error bars indicate standard error of the mean. p-value: *<0.05, Wilcoxon Signed-Rank Test. (f) Model depicting a conserved regulation of the piRNA pathway via interactions of the CCGTTA motifs with Ovo family TFs in animal ovaries and A-MYB in animal testes.

To test this hypothesis, we used mouse OVOL2 ChIP-seq data from female mouse primordial germ cell-like cells (PGCLCs; day 2 of induction) under transgenic expression of mouse *Ovol2* ^52^. We observed OVOL2 binding to CCGTTA motifs within or near promoters of the piRNA pathway genes (e.g., *Tdrkh*, *Tdrd7*, *Pld6*) (**Figure 7b and Supplementary Figure 7b**) which also showed upregulation in response to *Ovol2* expression in mouse ESCs or PGCLCs (**Supplementary Figures 7b and 7c**). To find out whether mouse OVOL2 also binds to ovary piRNA clusters, we defined ovary-specific, testis-specific, and shared piRNA clusters using small RNA-seq data from mouse ovaries and testes (>1 RPKM normalized counts mapping to the piRNA cluster regions; data from ^54^; coordinates obtained via proTRAC ^55^) and checked OVOL2 binding at these regions (**Figure 7c and Supplementary Figure 7a**). Our analysis revealed strong OVOL2 binding events to CCGTTA motifs within regions corresponding to ovary-specific piRNA clusters (**Figure 7c and Supplementary Figure 7a**). These OVOL2 binding events within motifs were found near ends or the centres of ovary piRNA clusters (**Figure 7a and Supplementary Figure 7a**). OVOL2 ChIP-seq peaks showed a significant enrichment at ovarian piRNA clusters but were absent from testis-specific piRNA clusters (**Figure 7a and Supplementary Figures 7a and 7d**).

Next, we asked whether there was evidence of human OVOL2 engagement at human ovarian piRNA clusters. We exploited small RNA-seq data from human fetal ovaries (data from ^56^) and analysed OVOL2 motif occurrences within human ovarian piRNA clusters (**Figures 7d**). Ranking of the human ovary piRNA clusters with small RNA-seq expression using data and coordinates defined in ^56^ (**Figure 7d and Supplementary Figure 7e**) revealed that the most highly expressed piRNA cluster in human fetal ovaries was cluster 71, which accounted for 54% of all piRNAs reads in ovaries (**Figures 7d and Supplementary Figure 7e**). The piRNA cluster 71 showed 2.5-fold enrichment for OVOL2 motifs over the genomic background (p<0.01, **Figure 7d**). The enriched OVOL2 motifs were highly concentrated near the 5’ end of cluster 71, suggesting a potential impact on promoter activity (**Figure 7d**). To determine if OVOL2 was physically associated with these motifs, we used OVOL2 ChIP-seq data from human induced pluripotent stem cells (iPSCs; WTC11, data from ENCODE ^57^). Indeed, OVOL2 ChIP-seq showed strong binding to the promoter region of the ovarian piRNA cluster 71 (**Supplementary Figure 7f**). Interestingly, piRNA cluster 14, the second highest ranking ovarian piRNA cluster, showed several strong OVOL2 binding regions across the cluster locus (**Supplementary Figure 7f**).

Similar to the binding pattern observed for mouse OVOL2 ChIP-seq, human OVOL2 ChIP-seq showed binding near promoters of the piRNA pathway genes (e.g., *TDRKH*, *TDRD7*, *TDRD3*, *PIWIL4*) when overexpressed in the human iPSCs (**Figure 7b and Supplementary Figure 7b**). Moreover, human OVOL2 binding near promoters of piRNA pathway genes (e.g., *TDRKH*, *PIWIL4*, and *TDRD7*) occurred at orthologous regions in the mouse genome where mouse OVOL2 showed corresponding binding events (**Figure 7b**). These orthologous OVOL2 binding events occurred at Ovo motifs that were highly conserved across the mammalian species based on multiple sequence alignments (**Figure 7b**). Overall, our results indicate that the Ovo TF family interactions with CCGTTA motifs in regulation of ovarian piRNA pathway components is a gene-regulatory feature that is conserved from flies to vertebrates.

### Ancient Ovo motifs are hallmarks of ovarian piRNA clusters in metazoans

To test whether engagement of Ovo to the ovary piRNA clusters is a general conserved feature of animal ovarian germ cells, we analysed Ovo motif enrichment within ovary piRNA clusters across multiple metazoan species. We characterized ovary piRNA clusters in species by their RPKM normalized counts (>1) mapping to the piRNA cluster regions (proTRAC piRNA cluster coordinates ^55,58^) using ovary small RNA-seq datasets from human (*Homo sapiens*), crab-eating macaque (*Macaca fascicularis*), mouse (*Mus musculus*), golden hamster (*Mesocricetus auratus*), cow (*Bos taurus*), zebrafish (Danio rerio), buff-tailed bumblebee (*Bombus terrestris*), African malaria mosquito (*Anopheles gambiae*), Arizona bark scorpion (*Centruroides sculpturatus*), and Pacific oyster (*Crassostrea gigas*). We then calculated the enrichments of OVOL2 motifs within the ovary piRNA clusters of each species. Our results revealed significant enrichment of OVOL2 motifs at ovary piRNA clusters in all ten species when compared to genomic backgrounds (p<0.05, Wilcoxon Signed-Rank Test, **Figure 7d**). Overall, our results suggest a conserved regulatory mechanism utilizing CCGTTA motifs underpinning the expression patterns of germline piRNA pathway in metazoan species, where Ovo/OVOL2 interaction with the motifs in ovaries serves as a female counterpart of the male-specific regulation by A-MYB in testes (**Figure 7e**).

## Discussion

Transcription factors and co-activators controlling the male germline piRNA pathway have been previously described in vertebrate testes ^31,59,60^; however, a female-specific counterpart of such a regulatory network controlling the expression of the female germline piRNA pathway in insect and vertebrate ovaries has remained an enigma. The identification of ovary-specific TFs that control the germline-specific piRNA pathway in female germ cells provides crucial insights into the regulation of transposon repression during oogenesis in animals.

In our study we revealed Ovo as the principal regulator of ∼70% of the germline-specific piRNA pathway genes in *Drosophila*. The transcription factor Ovo locus encodes both somatic and germline isoforms that are driven by distinct promoters and were once thought to be two distinct genes: the somatic *shavenbaby* (*svb*) and the germline *ovo* (isoforms *ovo-A* and *ovo-B*) (**Figure 5e**). The somatic *svb* is expressed only in embryonic, larval, and pupal epidermis cells, while the germline *ovo-B* is the major and the only essential isoform required for viability of female *Drosophila* germ cells ^53,61^. Our results indicate that enforced *ovo-B* expression activates the germline piRNA pathway components in the ovarian somatic cells (**Figure 4**) and *l(3)mbt* specifically represses the expression of *ovo-B* in the somatic cells by occupying the promoter region responsible for germline expression (**Figure 5e**).

Our results point towards a gene-regulatory model of Ovo in the female germline piRNA pathway in ovaries where transcription of germline piRNA pathway genes (e.g., *aub*, *ago3* and *vas*) could be directed by Ovo family TFs, analogous to the model previously suggested for A-MYB in the regulation of the male germline piRNA pathway in mice testes (**Figure 7f**) ^31^. Similar to A-MYB, Ovo and its homologs auto-regulate their own expression ^62–64^, and as our motif and ChIP-seq results reveal, show strong binding to germline piRNA pathway components. Our experiments show that Ovo is indeed able to activate expression of germline piRNA pathway components when ectopically expressed in fly somatic cells. This includes the nuage components Aub, Ago3 and Vas which are activated in expression in somatic cells in the presence of ectopic Ovo and assemble to form cellular structures resembling the nuage bodies of germ cells (**Figures 4d-f**). Our model is further supported by the multi-species analyses of OVOL2 ChIP-seq in mouse PGCLCs and human iPSCs revealing strong OVOL2 binding to the ovary piRNA clusters (**Figure 7**), thus indicating a high degree of conservation of this gene-regulatory mechanism across the animal kingdom. Previous work analysing ovarian piRNA clusters ^65^ reported an observation of strong A-MYB motif enrichments at ovarian piRNA clusters in macaques leading them to suggest a paradoxical involvement of the testis-specific A-MYB in driving transcription of ovarian clusters in ovaries; however, this enrichment is now compatible with our findings given that OVOL2 and A-MYB utilize the same motifs and these motifs would recruit OVOL2 in ovaries.

Ovo is well known for its conserved role in germline development in animals ranging from flies to mice ^53^. Germline development in flies and mammals follow distinct pathways ^66^. In flies, the preformation model states that the germline is established through maternally deposited germ cell determinants within oocytes while mammalian PGCs develop according to epigenetic/inductive mechanisms (epigenesis) where external cues dictate their developmental trajectory ^66,67^. Importantly, Ovo in flies is maternally deposited as a component of the germplasm and later pole cells, which establish the fly PGCs ^68^, while mammalian OVOL2 acts downstream of BMP signalling to control cell fate decisions during mammalian PGC specification in the epiblast ^53,69^, thus suggesting that Ovo family TFs control animal PGC development and germline piRNA pathway expression via both intrinsic and inductive mechanisms.

Interestingly, ∼36% of the mouse OVOL2 binding sites were also occupied by A-MYB in testes as both OVOL2 and A-MYB recognize the same core CCGTTA motif sequence (**Figure 7a**). This could be a mechanism driving the expression of the shared germline piRNA clusters in both ovaries and testes (**Supplementary Figure 7a**); however, other mechanisms such as motif affinity, chromatin accessibility or methylation must account for expression of sex-specific piRNA clusters. In our motif analysis, we noticed a pseudo-palindromic extension to the GTT core in the human A-MYB motif (**Figure 6a**) which could prefer A-MYB over OVOL2 at testis-specific clusters. Additionally, we observed that ∼52% of the OVOL2 binding events within the mouse ovary piRNA clusters occurred at the CpG islands (**Figure 7c and Supplementary Figure 7a**) which could be linked to sex-specific DNA methylation patterns in developing PGCs. TF binding sites are enriched at hypomethylated regions that evade the first wave of default *de novo* DNA-methylation ^70^, which starts from day 13 of embryonic development (E13.5) in male mouse PGCs ^71^ and coincides with the appearance of the pre-pachytene piRNAs ^54^; however, female PGCs undergo *de novo* methylation only later after birth ^72^. Temporal differences in DNA methylation and binding patterns by sex-specific TFs that direct female PGC development such as OVOL2 could therefore protect against methylation to control sex-specific piRNA cluster transcription. Of note, the mouse *Ovol2* gene encodes both the repressor isoform *Ovol2a* and the activator isoform *Ovol2b*. The binding patterns between OVOL2A and OVOL2B ChIP-seq were indistinguishable from each other at piRNA pathway components in mice, thus the mechanism driving the ovary-specific piRNA pathway could also depend on the interplay of these isoforms.

In fly ovaries, Ovo is continuously present in the nucleus of the germline cells and maternal Ovo persists in the embryo until zygotic Ovo is expressed; thus, Ovo binding sites could be potentially marking genomic locations important during the transition from one generation to the next ^61^. Ovo persists in ovarian germ cells and plays a role in female germline sex determination by controlling expression of Otu and Sxl ^73^. Our findings reveal its concurrent role in regulation of the female germline piRNA pathway. However, the testis-specific regulator controlling the male germline piRNA pathway in fly testes remains to be identified. The fly Myb TF is unlikely to act analogously to the vertebrate A-MYB, as it does not recognize the CCGTTA motifs (based on *de novo* motifs in ENCODE ChIP-seq data) and was reported to function as a weak repressor of piRNA factors in OSCs ^49^; therefore, a different testis-specific TF candidate concurrently controlling spermatogenesis and male piRNA pathway likely exists in fly testes.

Intriguingly, OVO-family TFs and MYB-family TFs belong to different TF lineages and harbour distinct DNA-binding domains yet are capable of binding and competing for the same genomic CCGTTA motifs ^64^, thus illustrating an example of convergent evolution where unrelated classes of DNA-binding domains evolve to bind to the same DNA elements. Under this model, the CCGTTA elements represent ancient evolutionary conserved *cis*-regulatory elements that interact with female germline-specific Ovo-family TFs in ovaries and male germline-specific A-MYB in testes of animals that in combination with co-factors such as TCFL5 ^60^ and other epigenetic mechanisms (e.g., chromatin accessibility, DNA-methylation, histone marks, L(3)mbt) control the germline piRNA pathway. Moreover, Ovo and Ovo-like TFs are comprised of both repressor and activator isoforms which could interplay to control the piRNA pathway in a sex- and tissue-specific manner. Overall, our results reveal a conserved gene-regulatory mechanism involving interactions of the ovary- and testis-specific TFs with the CCGTTA *cis*-regulatory elements behind the regulation of the germline piRNA pathway in animal ovaries and testes.

## Materials & Methods

### Cell culture and treatments

Wild-type (WT) OSCs ^74^ (DGRC Stock # 288), Δ*l(3)mbt*-OSCs ^34^ (DGRC Stock # 289), and fGS/OSS ^75^ (DGRC Stock # 191) were purchased from the *Drosophila* Genomics Resource Center (DGRC) and cultured at 26 °C in Shields and Sang M3 Insect Medium (Sigma-Aldrich, S3652-6X1L) supplemented with 10% fetal bovine serum (Sigma-Aldrich, F9665-500ML), 10% fly extract (DGRC Stock # 1645670), 0.6 mg/mL glutathione (Sigma-Aldrich, G6013-25G), and 10 mU/mL human insulin (Sigma-Aldrich, I9278-5ML). The fGS/OSS cells were cultured using 25% conditioned medium (from the old culture) and passaged before reaching 50% confluency.

### Fly stocks and handling

All flies were kept at 25°C on standard cornmeal food or propionic food. Control *w^1118^* flies were a gift from the University of Cambridge Department of Genetics Fly Facility. Transgenic *vas*-GFP fly strain (w[*]; TI{TI}vas[AID:EGFP]) was reported in ^76^ and obtained from Bloomington *Drosophila* Stock Center (#76126).

### Ovary dissections and cell dissociation

Flies were fed with yeast extract 2-3 days before the dissections. Ovaries were dissected into 1 mL ice-cold PBS and centrifuged at 500 g (4°C) for 5 min. Cells were dissociated using 5 mg/mL trypsin (Sigma-Aldrich, T1426-50MG) and 2.5 mg/mL collagenase-A (Sigma-Aldrich, 10103586001) in Ringer’s solution (600 µL for 148 ovaries) for 1 hr. at 800 rpm (30°C) in a thermomixer and passed through a 100μm cell strainer followed by addition of 400 µL of Schneider’s Medium (Thermo Fisher Scientific, 21720001) containing 10% FBS and centrifuged at 500 g (4°C) for 5 min. The supernatant was discarded and 600 µL of PBS was added to the pellet to wash by centrifugation at 500 g (4°C) for 5 min. The pellet was resuspended in 500 µL of PBS. Dissociation of ∼30 ovaries resulted in ∼500,000 cells.

### FACS sorting

The *w^1118^*strain flies and transgenic *vas*-GFP flies were fed with yeast extract 3 days prior to dissections. Ovaries from control *w^1118^* strain flies and transgenic *vas*-GFP flies were then dissected in ice-cold PBS and cells dissociated as described above. Cells were resuspended in PBS and sorted into GFP-positive and GFP-negative populations with FACS Aria cell sorter (BD Biosciences). Approximately ∼148 ovaries from dissections of ∼74 *vas*-GFP flies gave ∼36,000 Vas-GFP-positive cells after FACS sorting and 19 ovaries from the *w^1118^* strain flies were dissociated and used as controls.

### RNA isolation and RT-qPCR

RNA was isolated using RNeasy Mini Kit (QIAGEN, 74106) with RNase-Free DNase Set (QIAGEN, 79254) for DNase digestion during RNA purification as per manufacturer’s instruction. Reverse transcription was performed with 100 ng to 1 µg RNA using SuperScript™ IV Reverse Transcriptase (Thermo Fisher Scientific, 18090010). RT-qPCR was performed on 1:10 diluted cDNA using Fast SYBR™ Green Master Mix (Thermo Fisher Scientific, 4385610) with Bio-Rad C1000 Thermal Cycler. Primers were designed against exon-exon junctions for genes with introns and *rpL32* was used as an internal standard (**Supplementary Dataset 3**). Relative expression was analysed using the delta-delta Ct method ^77^.

### RNA-seq

For total RNA-seq, ribosomal depletion was performed using riboPOOLs for *Drosophila melanogaster* (Cambridge Bioscience) using 500 ng of total RNA as input. For mRNA-seq, poly(A) selection was performed using Poly(A) mRNA Magnetic Isolation Module (NEB, E7490) with 500 ng of total RNA as input. Libraries were prepared using The NEBNext® Ultra™ Directional RNA Library Prep Kit for Illumina (NEB, E7420L). Library quality was assessed using Agilent 2100 bioanalyzer (High Sensitivity DNA Kit). Libraries were quantified with KAPA Library Quantification Kit for Illumina (Kapa Biosystems, cat# KK4873) submitted to sequencing in NovaSeq 6000 System with 2 × 50 bp reads on SP flow cell.

### ATAC-seq

For cell lines, the omni-ATAC-seq protocol was used on 50,000 cells as described before ^78^. In brief, cells were lysed in ATAC-Resuspension Buffer (RSB) containing 0.1% NP40, 0.1% Tween20, and 0.01% Digitonin and washed out with cold ATAC-RSB containing 0.1% Tween-20. The transposition reaction was performed with 25 μl 2x TD buffer from and 2.5 μl transposase (100nM final) from Illumina Tagment DNA Enzyme and Buffer Small Kit (Illumina, 20034197), 16.5 μl PBS, 0.5 μl 1% digitonin, 0.5 μl 10% Tween-20, 5 μl H2O at 37°C for 30 minutes in a thermomixer with 1000 RPM mixing. For the whole-ovary ATAC-seq, the omni-ATAC-seq protocol was optimized by adjusting the numbers of ovaries used (6, 10, 20, 30) and tagmentation time (30 min to 1 hr). Zymo DNA Clean and Concentrator-5 Kit (Zymo, D4014) was used for clean-up and transposed fragments were amplified for 5 cycles [72 °C for 5 min, 98 °C for 30 s (98 °C for 10 s, 63 °C for 30 s, 72 °C for 1 min) × 5] using NEBNext 2xMasterMix (NEB, M0541S) with Ad1_noMX and v2_Ad2.* indexing primers followed by qPCR amplification to determine additional cycle numbers. Amplified DNA was purified using Zymo DNA Clean and Concentrator-5 Kit (Zymo, D4014) and eluted in 20 μl H_2_O. AMPure XP beads (Beckman Coulter, A63881) were used for double-sided bead purification. 100-600 bp fragments were selected using Blue Pippin (Sage Science, NC1025035). The library fragment size distribution was assessed using Agilent 2100 bioanalyzer on a High Sensitivity DNA Kit. Libraries were quantified with KAPA Library Quantification Kit for Illumina (Kapa Biosystems, KK4873) and submitted to sequencing in NovaSeq 6000 System with 2 × 50 bp reads on SP flow cell.

### ChIP-seq

For ectopic Ovo ChIP-seq, >30 x 10^6^ OSCs (n=2) were nucleofected with a *ovo-B-FLAG* construct using 0.5-0.8 μg per million cells, Nucleofector Kit V (Lonza, VVCA-1003), T-029 program on NucleofectorTM 2b Device (Lonza), and crosslinked 48 hours later by covering cells in 1% formaldehyde (FA) solution (50mM Hepes-KOH, 100mM NaCl, 1mM EDTA, 0.5mM EGTA, 1% FA) for 10 min at room temperature. Cell lysis was performed as described in ^79^. Sonication was performed with 15-18 cycles of 10 sec ON and 60 sec OFF (Fisherbrand™ Q705 Sonicator, 418-21 probe) in 3 mL of LB3 buffer (10 mM Tris–HCl, pH 8; 100 mM NaCl; 1 mM EDTA; 0.5 mM EGTA; 0.1% Na–Deoxycholate; 0.5% N-lauroylsarcosine; in 7 ml conical polypropylene tube) to shear DNA into 100-500 bp fragments. Chromatin immunoprecipitation was performed using 5-10 μg of primary mouse monoclonal ANTI-FLAG® M2 antibody (Merck, F1804-200UG) in 250 μL LB3 with 1% Triton X-100 overnight at 4 °C. Crosslinks were reversed by incubation at 65 °C for 16 h. Proteins and RNA were enzymatically digested. DNA purification was performed using phenol-chloroform extraction and ethanol precipitation. The purified ChIP DNA and 220 ng of whole cell extract DNA were used to prepare ChIP-seq libraries using NEBNext® Ultra™ II DNA Library Prep Kit for Illumina (NEB, E7645S) with PCR amplifications carried out for 16 cycles. The ChIP-seq libraries were assessed for fragment size distribution using Agilent 2100 bioanalyzer on a High Sensitivity DNA Kit, quantified with KAPA Library Quantification Kit for Illumina (Kapa Biosystems, KK4873), and sequenced on a NovaSeq 6000 System with 2 × 150 bp reads on SP flow cell.

### Overexpression screen

Candidate genes were amplified from *Drosophila melanogaster* ovary cDNA library using KOD Hot Start DNA Polymerase (Merck Millipore, 71086-4) and cloned into overexpression vectors under *act5* promoter with either N-terminal or C-terminal FLAG tags using Gibson Assembly® Master Mix (NEB, E2611L) at 50 °C for 1 hr. 9 μl of Mix & Go Competent Cells (Strain DH5 Alpha) were thawed on ice and transformed with 1 μl of diluted (2.5x in H_2_O) Gibson Assembly reaction by incubation for 5 min on ice before plating on LB plates containing the appropriate antibiotic. Colony PCR was performed to identify the colonies harbouring the ligated constructs followed by their inoculation into 3 mL broth containing 100 μg/mL ampicillin that were shaken overnight at 37 °C. The constructs were purified using QIAGEN Plasmid Plus Kits and Sanger sequenced to check for mutations and confirm the correct ligation. The constructs were then transfected into WT OSCs and Δ*l(3)mbt*-OSCs with Nucleofector Kit V (Lonza, VVCA-1003) using T-029 program and a NucleofectorTM 2b Device (Lonza). Cells were passaged 1 day prior to transfections and were 70-80% confluent at the time of transfections. Ectopic *ovo* overexpression vector carrying the FLAG-tagged *ovo-B* isoform matching NM_080338 transcript was used.

### siRNA knockdown experiments

Sense and antisense 21 nt siRNA sequences for target genes were designed using the DSIR tool (http://biodev.cea.fr/DSIR/DSIR.html) (**Supplementary Dataset 3**). The designed sequences were ordered from IDT and resuspended in 400 μl RNase-free water. Equal volumes of the resuspended sense and antisense siRNA were mixed then added to an equal amount of 2x annealing buffer (60 mM potassium acetate; 200 mM HEPES, pH 7.5). The mix was boiled for 5 min at 75 °C, then ramped down to 25 °C (−0.1 °C/second) to anneal the siRNA sequences into the siRNA duplex (100 μM final concentration =200 pmoles). 2 μL of the final 100 μM siRNA duplex was mixed with 100 μL of Nucleofection solution V (Lonza, VVCA-1003) and transfected into 10 million cells using a NucleofectorTM 2b Device and program T-029 (Lonza). Cells were plated into 12-well plates and the media was changed after 24 hr. RNA was either harvested at 48 hr or nucleofection was repeated after 48 hr and RNA was harvested at 96 hr.

### Western blots

Proteins were extracted from 1-3 x 10^6^ cells, washed in PBS (300 g, 5 min), and lysed in ice-cold RIPA buffer (Thermo Scientific, 89900) containing cOmplete™ Protease Inhibitor Cocktail EDTA-free tablet (Roche) (1 tablet per 20 mL of RIPA) under rocking condition at 4 °C for 30 min. The lysate was centrifuged 20,000g at 4 °C for 20 min and supernatant was transferred to a new tube. The protein concentrations were measured using Direct Detect® Spectrometer. 15 mg/mL of protein was used with NuPAGE 4X LDS Sample Buffer and NuPAGE™ 10X Sample Reducing Agent (Thermo Fisher Scientific) in 20 μL total volume and denatured at 70°C for 10 min followed by protein gel electrophoresis using NuPAGE™ 4-12%, Bis-Tris, 1.5mm 10 well, Mini Protein Gels and XCell SureLock Mini-Cell Electrophoresis System (Thermo Fisher Scientific) at 180V for 1hr at 4°C cold room. Precision Plus Protein™ All Blue Prestained Protein Standards (Bio-Rad, 1610373) was run in parallel as a ladder. Blotting was performed for 7 min with iBlot™ 2 Transfer Stacks, nitrocellulose, mini using iBlot™ 2 Gel Transfer Device. The nitrocellulose membrane was blocked for 1 hr at room temperature under gentle agitation using Intercept® blocking buffer (TBS) (LI-COR Biosciences). The membrane was rinsed with TBS buffer and incubated with the primary mouse monoclonal ANTI-FLAG® M2 antibody (Merck, F1804-200UG) (1:2,500) and the primary rabbit polyclonal anti-Tubulin (DM1A+DM1B) (Abcam, ab18251) (1:5,000) in TBS with 0.1% Tween 20 (TBST) overnight at 4 °C under gentle agitation. The membrane was washed 3 x 5-10 min with TBST and incubated with the secondary antibodies IRDye® 800CW Donkey anti-Mouse IgG Secondary Antibody (1:5,000) (LI-COR Biosciences, 926-32212) and IRDye® 680RD Donkey anti-Rabbit IgG Secondary Antibody (1:20,000) (LI-COR Biosciences, 926-68073) in TBST for 1 hr room temperature under gentle agitation. The membrane was washed 3 x 10 min with TBST and additional washed with TBS before visualizing with the Odyssey CLx Infrared Imaging System (LI-COR Biosciences).

### Immunofluorescence (IF) and microscopy

Coverslips were coated with fibronectin (1:50 in PBS) (Sigma-Aldrich, F0895-1MG) for 1 hr at 26 °C in 6-well plates and cells were plated on top. After 48 hr, cells attached to coverslips were fixed in 4% PFA in PBS (500 µL per well) for 15 min at room temperature and rinsed 3x with PBS. Cells were permeabilized with 0.2% Triton-X-100 in PBS for 10 min at room temperature and rinsed 3x with PBS. Blocking was performed with 0.1% Tween 20, 1% BSA (in PBS). Primary mouse monoclonal ANTI-FLAG® M2 antibody (Merck, F1804-200UG) (1:500), primary rabbit monoclonal DYKDDDDK Tag (D6W5B) ANTI-FLAG antibody (Cell Signalling, 14793S), primary rabbit polyclonal anti-Piwi (1:500, AB1159, in-house, lot# PUC5368), primary mouse monoclonal anti-Aub (1:500, Brennecke lab ^80^, 8A3-D7), primary rabbit polyclonal anti-Aub (Brennecke lab ^3^), primary mouse monoclonal anti-Ago3 (1:500, Brennecke lab, 7B4-C2), primary rabbit polyclonal anti-Ago3 (Brennecke lab ^3^), primary mouse monoclonal anti-Yb (Siomi lab, 8H12B12, 180803), primary rat monoclonal anti-Vasa (DSHB, AB_760351) and primary mouse monoclonal anti-Lamin Dm0 (DSHB, adl67.10) antibodies were diluted in 0.1% Tween 20, 0.2% BSA, 1X PBS (500 µL/well) and added onto coverslips in each well. Incubation with primary antibodies was performed overnight at 4 °C in the cold room under gentle agitation. Coverslips were washed 3x for 5 min with PBS (0.1% Tween 20) under gentle agitation. Goat anti-Mouse IgG (H+L) Highly Cross-Adsorbed Secondary Antibody, Alexa Fluor 647 (Thermo Fisher Scientific, A-21236) and Goat anti-Rabbit IgG (H+L) Cross-Adsorbed Secondary Antibody, Alexa Fluor™ 488 (Thermo Fisher Scientific, A-11008) were diluted 1:500 in 0.1% Tween 20, 0.2% BSA, 1X PBS and added on coverslips (500 µL/well) for 1 hr at room temperature under gentle agitation (covered with aluminium foil) followed by 3x 5 min washes with PBS (0.1% Tween 20). Coverslips were incubated with DAPI at 1:10,000 dilution in PBS (0.1% Tween 20) for 10 min at room temperature (covered with aluminium foil) followed by 2x 5 min washes with PBS (0.1% Tween 20) under gentle agitation. Coverslips were mounted on glass slides using a drop of ProLong™ Diamond Antifade Mountant (Thermo Fisher Scientific, P36961). Microscopy was performed with the Confocal Laser Scanning System Leica SP8 and image analysis was performed using Leica Application Suite X (v3.5.7.23225) and Huygens Professional (v20.04).

### ATAC-seq data analysis

The read quality was assessed with FastQC (v0.11.8). The paired-end reads were trimmed of adapter sequences TCGTCGGCAGCGTCAGATGTGTATAAGAGACAG and GTCTCGTGGGCTCGGAGATGTGTATAAGAGACAG using Cutadapt tool (v1.18; default parameters) ^81^. Burrows-Wheeler Aligner (BWA) (v0.7.17, bwa mem -M -t 4) ^82^ was used to align the trimmed paired reads to 2014 (BDGP Release 6 + ISO1 MT/dm6) assembly of the *D. melanogaster* genome. Duplicates were marked using Picard tool (v2.9.0) (MarkDuplicates, validation stringency=lenient). SAMtools (v1.9) was used for indexing and filtering ^83^. The quality metrics for the aligned ATAC-seq reads were assessed using ataqv (v1.0.0) (https://github.com/ParkerLab/ataqv) ^84^. The ATAC-seq peaks were called with MACS2 (v2.1.1.20160309) using --nomodel -- shift -37 --extsize 73 parameters and FDR cut-off q ≤ 0.05 ^85^. Differential accessibility analysis was performed using DiffBind package (v3.0.15) ^86^. RPKM normalized bigWig files were generated using deepTools (v3.5.1, bamCoverage) ^87^. Peak intersections were performed using bedtools (v2.30.0) ^88^ ATAC-seq heatmaps and profiles were generated using plotHeatmap and plotProfile tools in the deepTools (v3.5.1). Genome browser visualizations were done using the UCSC Genome Browser ^89^.

### RNA-seq data analysis

The reads were trimmed of adapter sequences using Cutadapt tool (v1.18; -m 1 specified to not have reads trimmed to zero) ^81^. The trimmed reads were aligned to the genome assemblies using the RNA-seq aligner STAR (v2.7.3a) ^90^. Gene counts were calculated with the featureCounts tool (Subread package v1.5.3; -s 2 for stranded libraries prepared by dUTP method) ^91^. *D. mel* gene annotations were taken from Drosophila_melanogaster.BDGP6.28.100.gtf in the Ensembl genome database ^92^. Samtools (v1.9) was used to merge bam files from replicates ^83^. Differential gene expression analysis was performed using DESeq2 (v1.30.1) ^93^. The RPKM normalized bigWig files were generated for each strand using bamCoverage tool (-- filterRNAstrand specified for dUTP stranded libraries) in deepTools (v3.5.1) ^87^. Raw data for publicly available RNA-seq reads were downloaded from sources listed in **Supplementary Dataset 3** and processed as above. Genome browser visualizations were done using the UCSC Genome Browser ^89^.

### Single-cell RNA-seq analysis

The single-cell RNA-seq data matrices were downloaded from sources listed in **Supplementary Dataset 3** and clustered using Seurat 4.0 toolkit (v4.3.0.1) ^94^. Normalizations were performed with SCTransform() function. Differential expression analysis was performed using FindMarkers() function. Pseudotime analysis was done with Monocle 3 tool (v3.0) ^95^. Correlations were done with the Pearson correlation coefficients and p-values were corrected for multiple testing using Bonferroni correction. The custom code for single-cell RNA-seq co-expression analysis is available at GitHub (https://github.com/alizadaa/Single-cell_RNA-seq_Co-expression_Analysis_Ovary).

### ChIP-seq data analysis

The reads were trimmed of adapter sequences using Cutadapt tool (v1.18) ^81^. The trimmed reads were aligned to the genome assemblies using Burrows-Wheeler Aligner (BWA) (v0.7.17, bwa aln) ^82^. SAMtools (v1.9) was used for sorting, merging, and indexing ^83^. ChIP-seq peaks were called using MACS3 (v3.0.0a6) callpeak command (-q 0.01) ^85^ with data from ChIP input (the whole cell lysate) used as the control (-c). The reproducibility of ChIP-seq peaks between replicates was evaluated using the irreproducible discovery rate (IDR) tool (v2.0.2) ^96^. The RPKM normalized bigWig files were generated using bamCoverage tool in deepTools (v3.5.1) ^87^ with the parameter (--extendReads 120) specified. The input-normalized bigWig files were generated using the bamCompare tool in the deepTools (v3.5.1) ^87^. Signals from bigWig files were quantified using the multiBigwigSummary tool in deepTools (v3.5.1) ^87^. ChIP-seq heatmaps and profiles were generated using the plotHeatmap and plotProfile tools in the deepTools (v3.5.1) ^87^. Raw data for publicly available ChIP-seq datasets were downloaded from sources listed in **Supplementary Dataset 3** and processed as describe above. Genome browser visualizations were done using the UCSC Genome Browser ^89^.

### Small-RNA-seq and piRNA cluster analysis

The raw small RNA-seq reads from ovaries and testes of human (*Homo sapiens*), crab-eating macaque (*Macaca fascicularis*), mouse (*Mus musculus*), golden hamster (*Mesocricetus auratus*), cow (*Bos taurus*), zebrafish (Danio rerio), buff-tailed bumblebee (*Bombus terrestris*), African malaria mosquito (*Anopheles gambiae*), Arizona bark scorpion (*Centruroides sculpturatus*), and Pacific oyster (*Crassostrea gigas*) were downloaded from sources listed in **Supplementary Dataset 3**. The reads were trimmed of adapter sequences using Cutadapt tool (v1.18) ^81^ and aligned to the respective genome assemblies using the RNA-seq aligner STAR (v2.7.3a) ^90^. Samtools (v1.9) was used to merge bam files from replicates ^83^. The RPKM normalized bigWig files were generated using bamCoverage tool in deepTools (v3.5.1) ^87^. The piRNA cluster coordinates were taken from the piRNA cluster database (https://www.smallrnagroup.uni-mainz.de/piRNAclusterDB/) ^58^. Ovary and testis piRNA clusters were defined and ranked using the RPKM normalized counts (>1) mapping to the piRNA cluster regions as calculated from bigWig files of the ovary and testis small RNA-seq datasets using multiBigwigSummary tool in deepTools (v3.5.1)^87^. Genome browser visualizations were done using the UCSC Genome Browser ^89^.

### Motif analysis, peak enrichments, and clustering

Motif scanning was performed using FIMO tool in MEME Suite (v5.4.1) (https://meme-suite.org/meme/index.html) ^97^. *De novo* ChIP-seq motifs were generated using MEME-ChIP tool in MEME Suite (v5.4.1) ^98^. List of *Drosophila* motifs were downloaded from FlyFactorSurvey (http://pgfe.umassmed.edu/TFDBS/) database ^99^. ^100^fastaFromBed (v2.26.0) was used for conversion of bed file coordinates into fasta format using *D.mel* BDGP6.28.dna.toplevel genome. Data was visualized with UCSC genome browser ^89^. ChIPseeker (v1.36.0) ^101^ was used to annotate the genomic features and distances based on TxDb.Dmelanogaster.UCSC.dm6.ensGene annotation package. The radial tree clustering of human TF motifs was adapted from the JASPAR database ^102^ (https://jaspar.elixir.no/matrix-clusters/) (JASPAR 2022 vertebrates CORE; RSAT - matrix-clustering ^103^). Orthologous regions between the species genomes were derived using the UCSC LiftOver tool (https://genome.ucsc.edu/cgi-bin/hgLiftOver) ^104^. The species classification tree was generated using the NCBI taxonomy database ^105^ and visualized in TreeViewer ^106^. Multiple sequences alignments of motifs and phylogenetic trees were based on Multiz alignments (27-way insect alignment and 46-way vertebrates) ^107^ from the UCSC conservation tracks ^108–110^.

## Data availability

RNA-seq, ATAC-seq and ChIP-seq data generated in this study have been deposited to GEO (accession number pending).

## Code availability

Custom code is available at GitHub (https://github.com/alizadaa/Single-cell_RNA-seq_Co-expression_Analysis_Ovary) ^111^.

## Acknowledgements

We thank Anna Sobieszek for preparing RNA-seq libraries from OSC knockdown samples. We thank Hannon group members for fruitful discussions and Susanne Bornelöv for advice on computational analyses. We thank the Scientific Computing core at the CRUK Cambridge Institute for HPC resources, the Flow Cytometry core for FACS services, Research Instrumentation and Cell Services for liquid nitrogen storage and mycoplasma testing, the Light Microscopy core for training and advise, and the Genomics core for sequencing services. GJH is a Royal Society Wolfson Research Professor (RSRP\R\200001). This research was funded in whole, or in part, by Cancer Research UK (G101107) and the Wellcome Trust (110161/Z/15/Z and 226627/Z/22/Z).

## Author contributions

AA, BCN and GJH conceived the study. AA and BCN designed the experiments and interpreted the results. AA performed all wet-lab experiments and computational analyses and wrote the first manuscript draft. BCN and GJH supervised the project and wrote the manuscript. All authors read and approved the final version.

## Source data

**Supplementary Dataset 1.** Single-cell RNA-seq correlation scores and adjusted p-values from the co-expression analysis.

**Supplementary Dataset 2.** Differential gene expression changes between ovarian cell types and under enforced Ovo expression.

**Supplementary Dataset 3.** Data sources for genomics analyses. siRNA and RT-qPCR sequences.

## Supplementary Figures and Legends

**Supplementary Figure 1.**
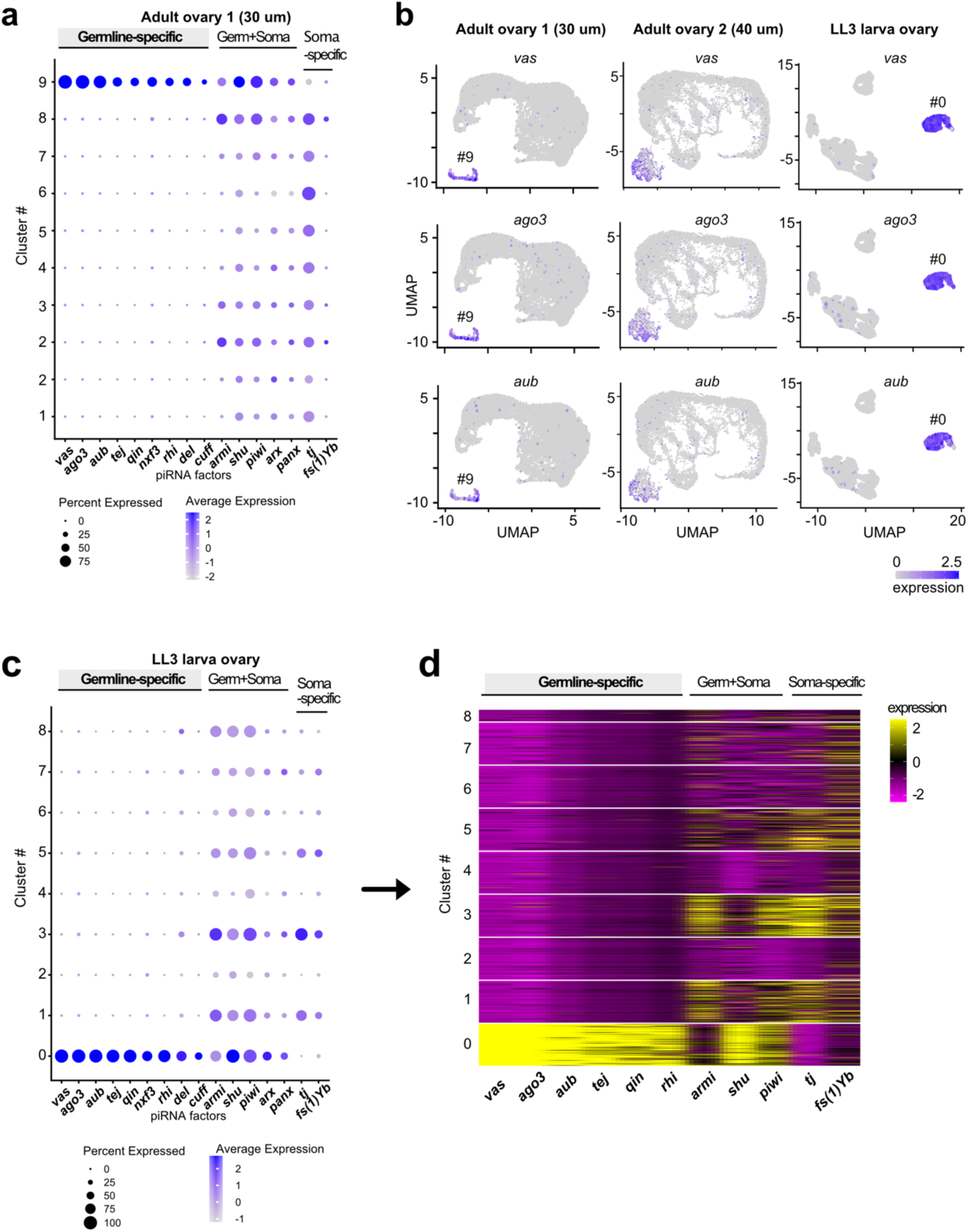
*Drosophila* ovary single-cell RNA-seq datasets reveal specific expression of the germline piRNA pathway genes in the ovarian germ cell clusters. (a-b) Dot plot and feature plots showing the expression of the germline-specific, shared, and soma-specific piRNA pathway genes across the cell clusters identified from the adult ovary dataset 1 (30 µm cell strainer; cluster 9 is the germline cluster; data from ^35^) and (b-d) across the clusters identified from the late third-stage larva (LL3) ovary (40 µm cell strainer; cluster 0 is the germline cluster; data from ^37^).

**Supplementary Figure 2.**
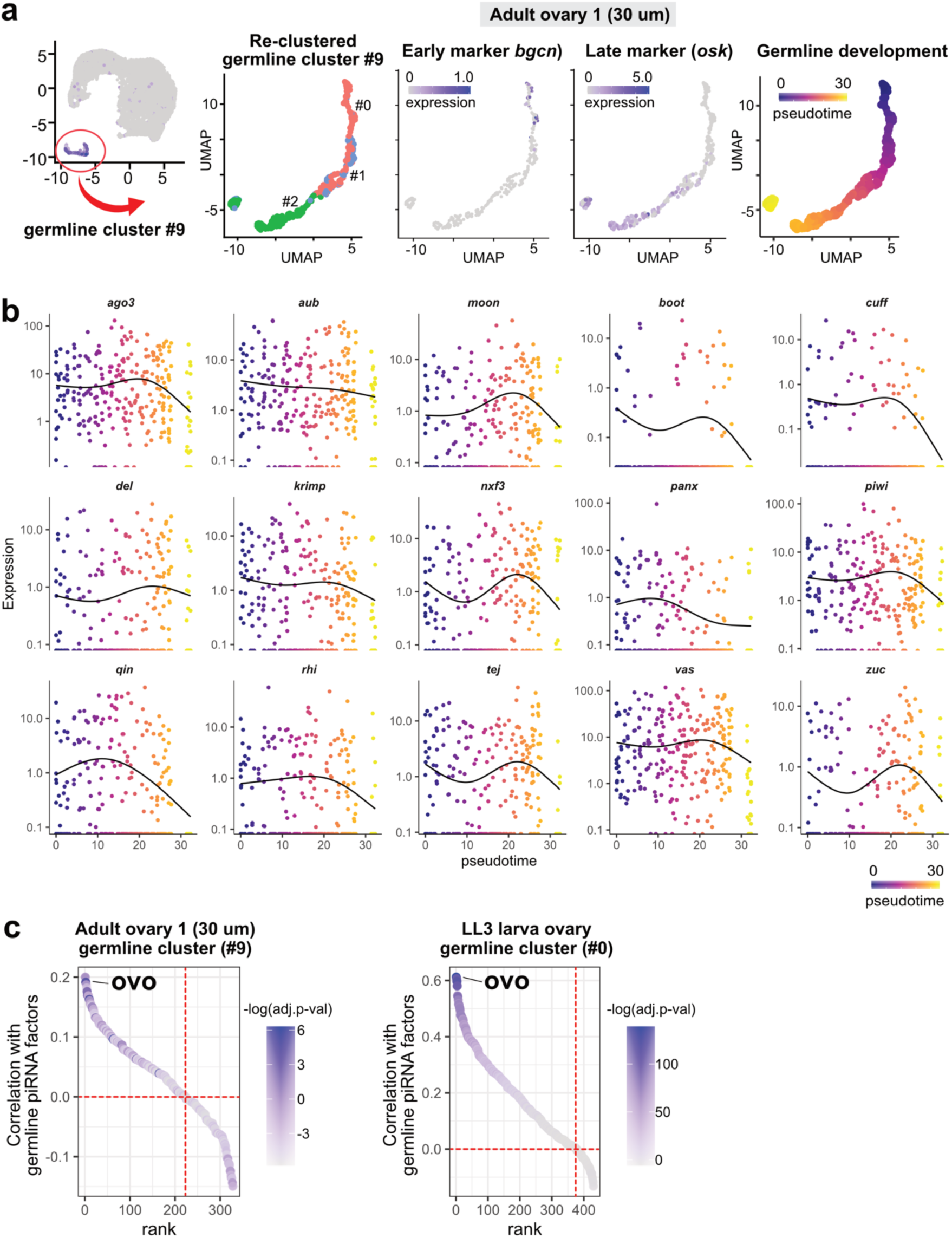
Ovo is the top transcription factor co-expressed with germline piRNA pathway genes throughout germ cell stages. (a) UMAP clustering showing the re-clustered germline cluster 9 from the adult ovary 1 dataset and pseudotime trajectory of germline development computed by rooting the early *bgcn*-expressing germline stem cells (GSCs) as the starting point. (b) The expression pattern of the piRNA pathway genes along the pseudotime trajectory of the germline development in the adult ovary 1 dataset. (c) Ranking of the DNA-binding transcription factors (TFs) by the average expression correlation (Pearson’s r) with the germline piRNA pathway genes *aub*, *vas*, *qin*, and *ago3* in the re-clustered germline cluster 9 of the adult ovary 1 (left) and the re-clustered germline cluster 0 of the larva ovary (right). The colour scales show correlation p-values adjusted with Bonferroni correction for multiple testing.

**Supplementary Figure 3.**
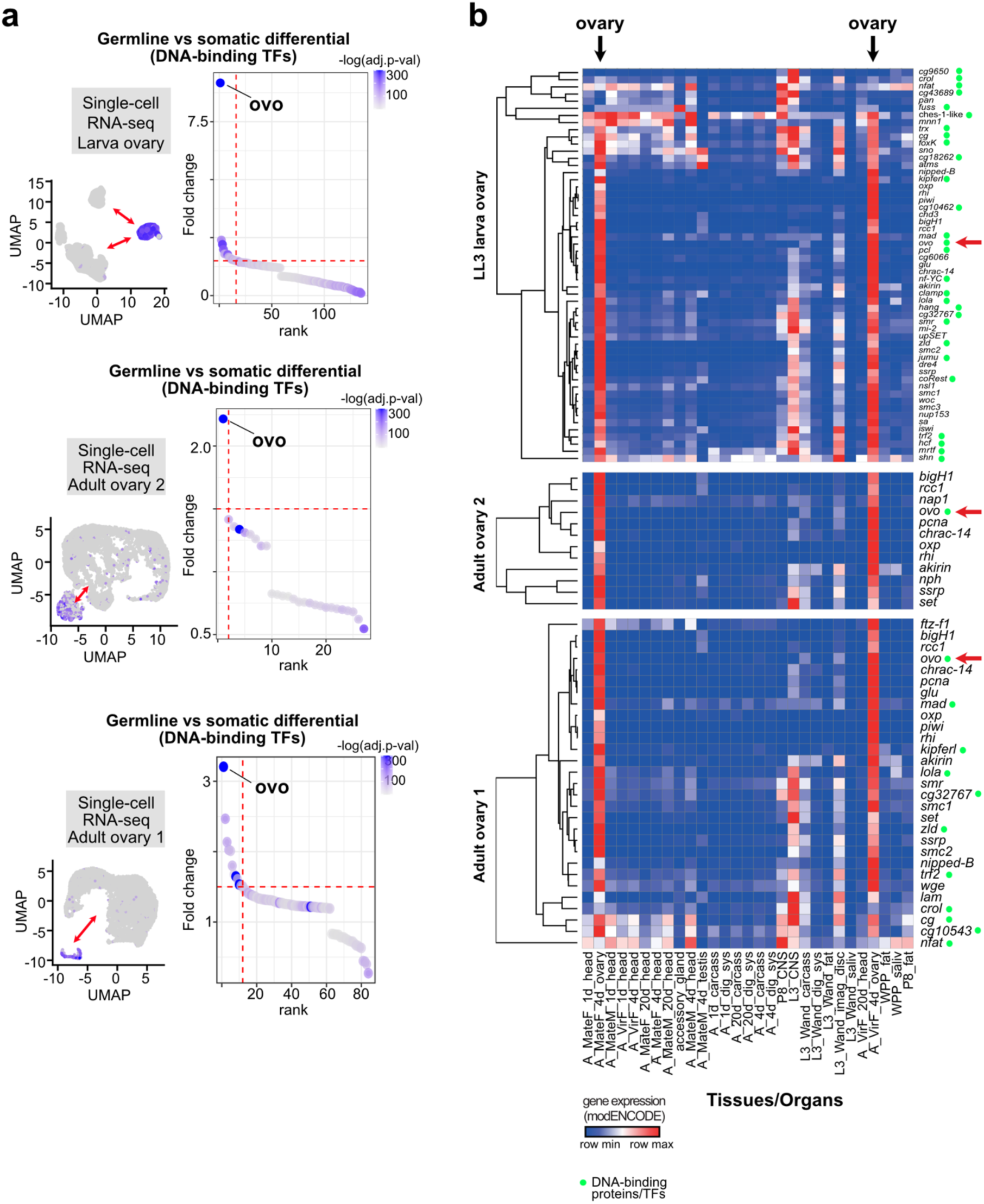
Ovo is the major ovarian germline transcription factor in the *Drosophila*. (a) Differential gene expression (Seurat package, FindMarkers tool) between the germline and somatic clusters of the LL3 larva and adult ovary single-cell RNA-seq datasets from ^35–37^ showing the top-ranking germline-enriched DNA-binding TFs. The colour scales show the non-parametric Wilcoxon rank sum test p-values adjusted with Bonferroni correction for multiple testing. (b) Cross-tissue expression (bulk RNA-seq, modENCODE) of the top-ranking chromatin-binding and DNA-binding candidate genes showing the highest co-expression with the germline piRNA factors in ovary single-cell RNA-seq datasets. Red arrows highlight *ovo* expression in ovaries.

**Supplementary Figure 4.**
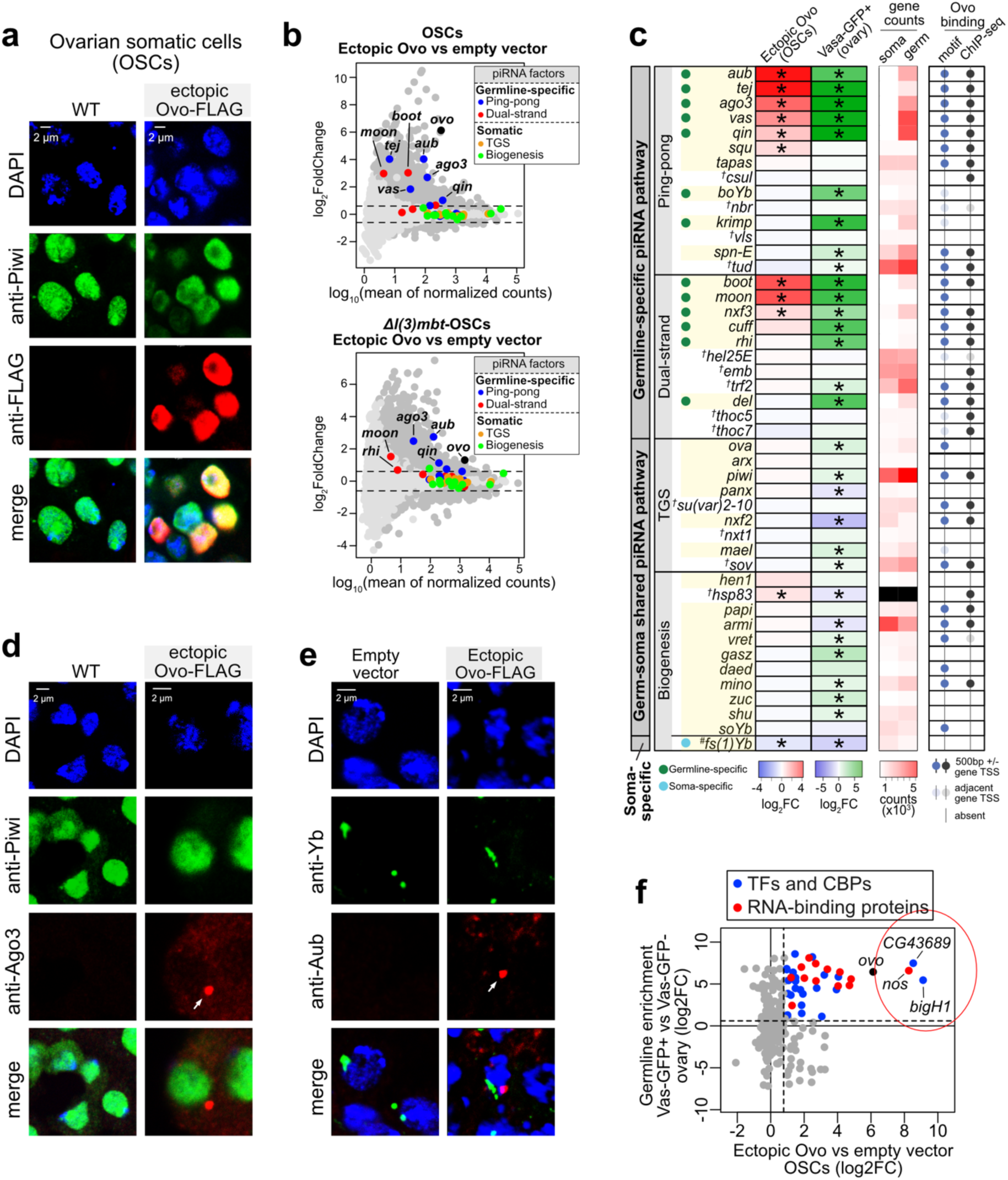
Enforced expression of *ovo-B* in ovarian somatic cells (OSCs) activates the expression of the germline-specific piRNA pathway components. (a) Immunofluorescence images showing the nuclear localization of the Ovo-FLAG protein in OSCs after nucleofection with the *ovo-FLAG* construct (48-72 hr; Ovo-B isoform, NM_080338 transcript; DAPI indicates the nuclear DNA and Piwi indicates the nucleoplasm). (b) MA-plots showing the fold-changes and normalized counts of gene expression (Deseq2, RNA-seq n=3 replicates from distinct samples) for all genes following the nucleofection of the wild-type (WT) OSCs (top) and Δ*l(3)mbt* OSCs (bottom) with the *ovo-FLAG* construct (Ectopic Ovo) relative to the nucleofection with an empty vector. Data point colours: Light grey =not significant (Deseq2 adjusted p-value >0.1); Dark grey= significant (Deseq2 adjusted p-value <0.1). The piRNA pathway genes labelled according to the colour key (top right corners). TGS =transcriptional gene silencing. (c) Table summarising the fold-changes in gene expression for all the genes involved in the piRNA pathway (germline and somatic; TGS= transcriptional gene silencing) following the nucleofection with the *ovo-FLAG* construct) relative to the nucleofection with an empty vector in OSCs, and the fold-enrichments of the same genes in the Vas-GFP+ germline cells compared to the Vas-GFP- somatic cells from the FACS-sorted transgenic Vas-GFP ovaries (Deseq2, RNA-seq n=3 replicates from distinct samples). The piRNA pathway-specific genes are highlighted in yellow; †=ubiquitous genes that are not specific to the piRNA pathway; # =somatic-specific piRNA pathway genes; *<0.01, adjusted p-values from Deseq2; normalized gene counts (Deseq2) are shown for Vas-GFP- (soma) and Vas-GFP+ (germ) cells (*hsp83* in black is excluded due to high expression levels). The presence of Ovo motifs and Ovo ChIP-seq peaks (ENCODE) within 500 bp +\- of gene TSS is shown with dots on the right side of the table. Transparent dots represent non-specific motifs and peaks within 500 bp +\- of gene TSS that are closer to a different/neighbouring gene. (d) Immunofluorescence images showing the presence of Ago3 proteins as peri-nuclear nuage-like foci (arrowheads) in the OSCs nucleofected with the ovo-FLAG construct (DAPI indicates the nuclear DNA and Piwi indicates the nucleoplasm). (e) Immunofluorescence images showing distinct localization of the nuage-like foci (red, Aub) appearing within the *ovo-FLAG* nucleofected OSCs from the somatic Yb bodies (green) normally present in OSCs. (f) Plot showing the Ovo target genes (x-axis; log_2_ fold-change response to ectopic Ovo expression in OSCs) that are germline-enriched (y-axis; vas-GFP+) and have gene-regulatory functions (TF=transcription factor, CBP=chromatin binding protein).

**Supplementary Figure 5.**
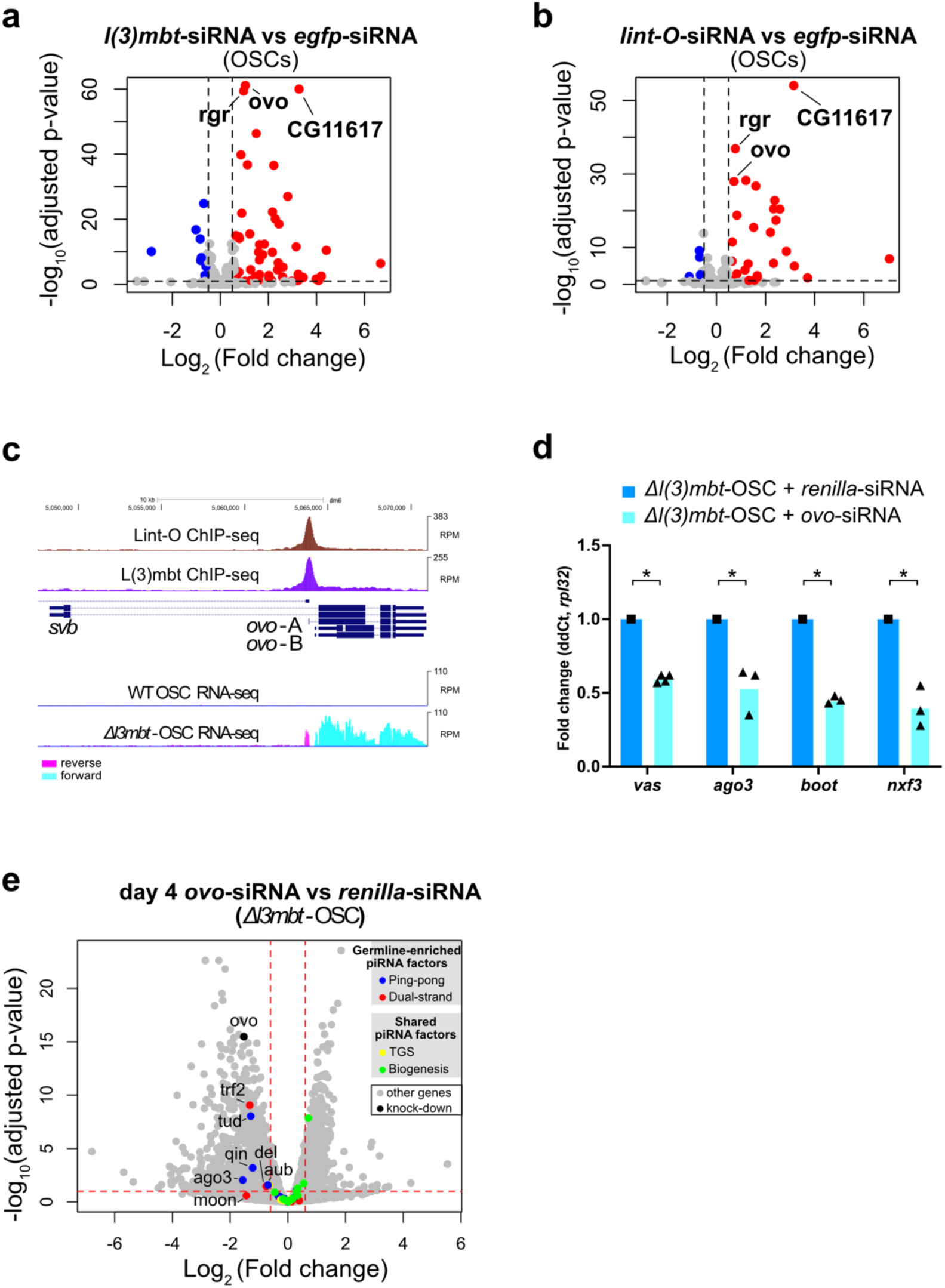
The complex of L(3)mbt with Lint-O indirectly represses the expression of the germline piRNA pathway genes in ovarian somatic cells (OSCs) via inhibition of germline *ovo-B* transcription. (a) siRNA knock-downs of *l(3)mbt* and (b) *lint-O* in ovarian somatic cells (OSCs) result in significant upregulation of *ovo*. (Deseq2; RNA-seq; n=3 replicates from distinct samples; data from ^49^) (c) Lint-O ChIP-seq showing a strong binding event at the germline *ovo* promoter (Ovo-B isoform) in OSCs where L(3)mbt ChIP-seq also shows a strong binding signal (rpm; merged n=2 replicates from distinct samples, data from ^49^). (d) *ovo* siRNA knock-down experiments in *Δl(3)mbt* OSCs (RT-qPCR; n≥3 replicates from distinct samples; p-value: *<0.01, one-tailed two-sample t-test). (e) Volcano plot showing downregulation of the germline-specific piRNA pathway genes on day 4 of *ovo* siRNA knock-downs in *Δl(3)mbt* OSCs using differential RNA-seq analysis (Deseq2) between *ovo* siRNA and *renilla* siRNA knock-downs (n=3 replicates from distinct samples).

**Supplementary Figure 6.**
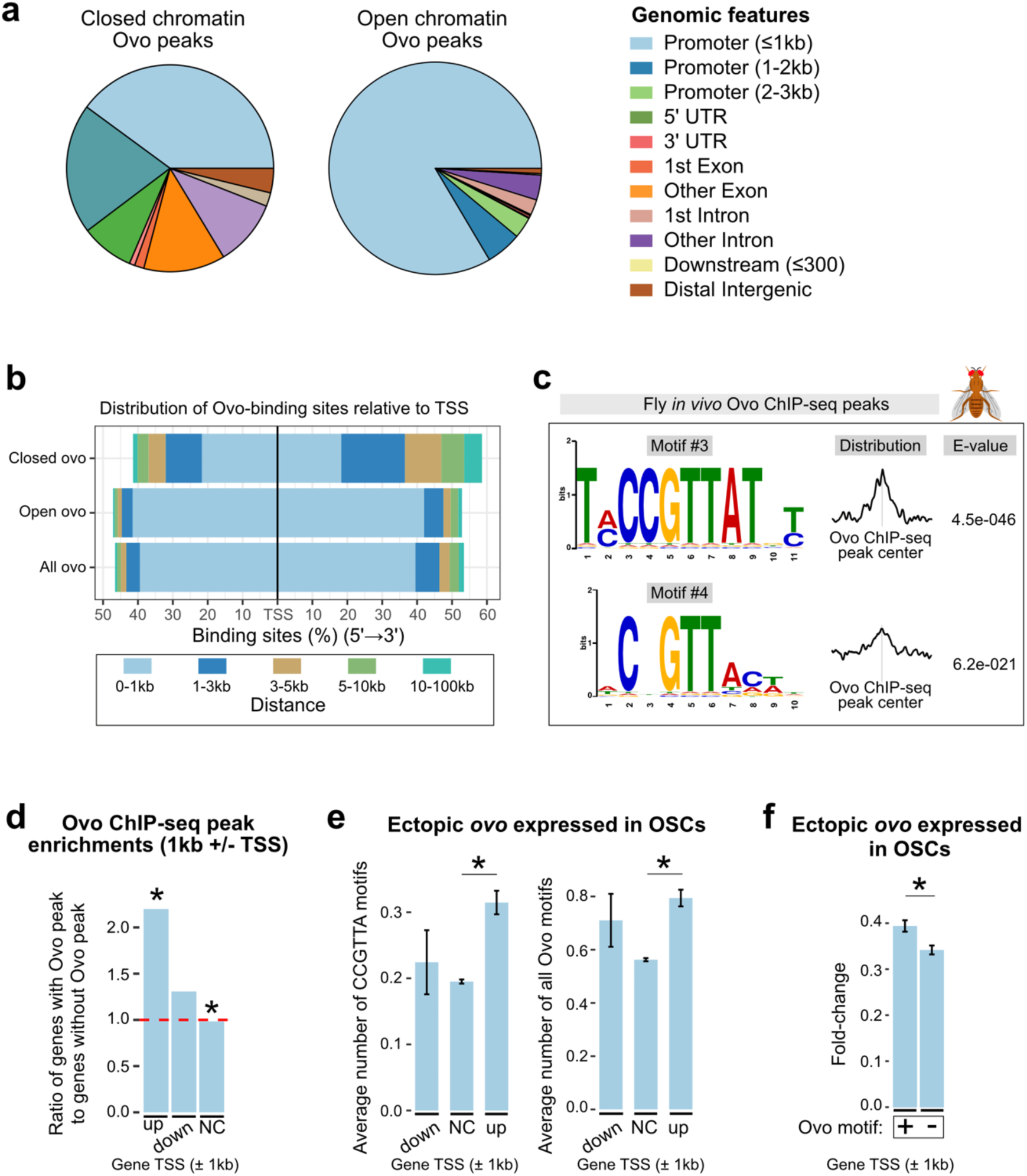
Genomic features of Ovo binding sites in *Drosophila*. (a) Genomic annotations (UCSC; Ensembl genes; dm6) of Ovo ChIP-seq peaks (data from ENCODE, whole fly) corresponding to open and closed chromatin states in fly ovaries based on ovary ATAC-seq peaks using ChIPseeker. (b) Distribution of open (ATAC-seq positive) and closed (ATAC-seq negative) Ovo ChIP-seq peaks relative to the gene transcription start sites (TSS). (c) The 3rd and 4th top-scoring *de novo* motifs discovered within fly Ovo ChIP-seq peaks using MEME-ChIP. (d) Enrichment of Ovo ChIP-seq peaks within 1kb ± TSS of the genes that are upregulated, downregulated or do not change in expression in response to the ectopic Ovo expression in OSCs (p-value: *<0.01, Fisher’s exact test). (e) Average number of CCGTTA Ovo motifs (B1H-derived; from FlyFactorSurvey database) and the average number of all Ovo motifs (both CCGTTA and CNGTTA motifs) within 1kb ± of TSS of genes that were downregulated (down), unresponsive (NC=no change), and upregulated (up) in response to the ectopic Ovo expression in OSCs. (f) Average fold-changes in response to the ectopic Ovo expression in OSCs compared between the genes that harbour (+) and lack (-) Ovo motif within 1kb ± of their TSS. Error bars indicate standard error of the mean. p-value: *<0.01, one-tailed two-sample t-test.

**Supplementary Figure 7.**
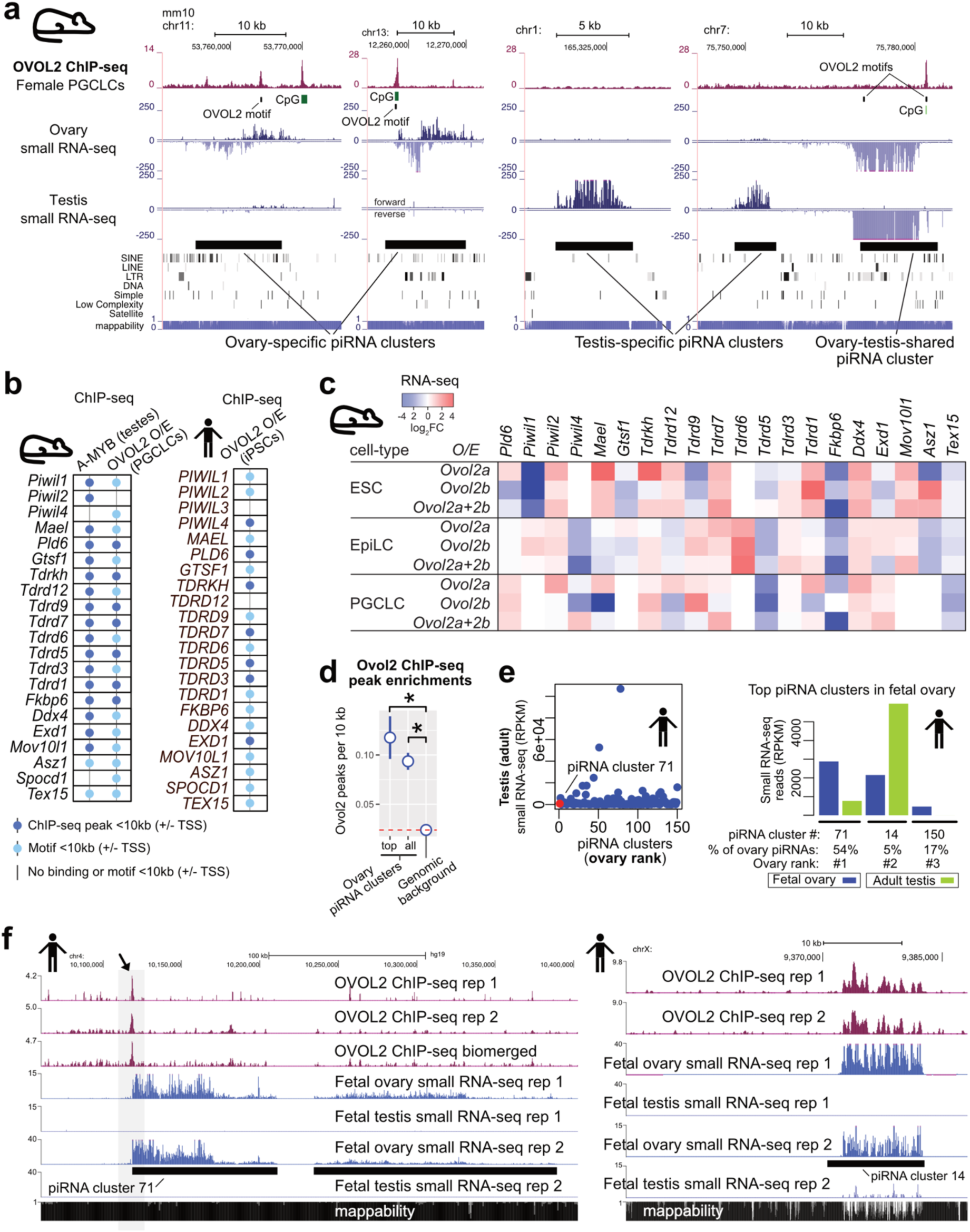
Human and mouse OVOL2 binding to ovarian piRNA clusters. (a) OVOL2 ChIP-seq (rpm; isoform 2A; merged n=2 replicates from distinct samples) from female mouse primordial germ cell-like cells (PGCLCs; day 2 of PGCLC induction; overexpressing transgenic mouse *Ovol2a*; data from ^52^) showing OVOL2 binding events at the mouse ovary piRNA clusters (small RNA-seq data from ^54^). ChIP-seq data for OVOL2A isoform is shown. The OVOL2B isoform shows the exact same binding patterns as OVOL2A. (b) Table summarizing binding of mouse A-MYB to piRNA factors in mouse testes (ChIP-seq data n=1 from ^31^), binding of mouse OVOL2 to piRNA factors in female mouse PGCLCs overexpressing (O/E) transgenic mouse *Ovol2a* (day 2 of PGCLC induction, OVOL2A isoform ChIP-seq data from ^52^; merged n=2 replicates from distinct samples), and binding of human OVOL2 to piRNA factors in human induced pluripotent cell (iPSC) line WTC11 (male) overexpressing (O/E) human *OVOL2* (ChIP-seq data from ENCODE; merged n=2 replicates from distinct samples). (c) Changes in expression of mouse piRNA pathway genes (log_2_ Fold change) under transgenic expression of mouse *Ovol2a* and *Ovol2b* throughout the *in vitro* differentiation of mouse embryonic stem cells (ES cells) to PGCLCs (day 2 of PGCLC induction) via epiblast-like cells (EpiLCs) (n=2 replicates from distinct samples, RNA-seq data from ^52^). (d) Numbers of mouse OVOL2 ChIP-seq peaks (isoform 2A, merged n=2 replicates from distinct samples; data from ^52^) per 10 kb at the piRNA clusters compared to the genomic background (all genomic regions that are not piRNA clusters). top= top-expressed ovary piRNA clusters (>20 rpm), all= all ovary piRNA clusters (>1 rpm). Error bars indicate standard error of the mean. p-value: *<0.05, Wilcoxon Signed-Rank Test. (e) The expression levels of the human piRNA clusters in the human adult testes ranked by their expression levels in human fetal ovaries (rpkm; data from ^56^). Barplots (right) are showing the expression levels of the top 3 highest expressed human fetal ovary piRNA clusters (#71, #14 and #150) in fetal ovaries and adult testes (rpkm). % of ovary piRNAs indicates the percentage of all fetal ovary piRNA reads mapping to the indicated clusters (data from ^56^). (f) OVOL2 ChIP-seq from human induced pluripotent cell (iPSC) line WTC11 overexpressing (O/E) ectopic OVOL2 construct (data from ENCODE) showing OVOL2 binding to the top expressed human fetal ovary piRNA clusters #71 (top) and #14 (bottom). Fetal ovary small-RNA-seq and fetal testes small-RNA-seq tracks are shown below (data from ^56^). Mappability tracks from Duke Uniq 35 are shown below.

